# Decoding RNA Metabolism by RNA-linked CRISPR Screening in Human Cells

**DOI:** 10.1101/2024.07.25.605204

**Authors:** Patrick J. Nugent, Heungwon Park, Cynthia L. Wladyka, Katharine Y. Chen, Christine Bynum, Grace Quarterman, Andrew C. Hsieh, Arvind Rasi Subramaniam

## Abstract

RNAs undergo a complex choreography of metabolic processes in human cells that are regulated by thousands of RNA-associated proteins. While the effects of individual RNA-associated proteins on RNA metabolism have been extensively characterized, the full complement of regulators for most RNA metabolic events remain unknown. Here we present a massively parallel RNA-linked CRISPR (ReLiC) screening approach to measure the responses of diverse RNA metabolic events to knockout of 2,092 human genes encoding all known RNA-associated proteins. ReLiC screens highlight modular interactions between gene networks regulating splicing, translation, and decay of mRNAs. When combined with biochemical fractionation of polysomes, ReLiC reveals striking pathway-specific coupling between growth fitness and mRNA translation. Perturbing different components of the translation and proteostasis machineries have distinct effects on ribosome occupancy, while perturbing mRNA transcription leaves ribosome occupancy largely intact. Isoform-selective ReLiC screens capture differential regulation of intron retention and exon skipping by SF3b complex subunits. Chemogenomic screens using ReLiC decipher translational regulators upstream of mRNA decay and uncover a role for the ribosome collision sensor GCN1 during treatment with the anti-leukemic drug homoharringtonine. Our work demonstrates ReLiC as a versatile platform for discovering and dissecting regulatory principles of human RNA metabolism.

## Introduction

RNAs are carriers of genetic information, scaffolds for protein complexes, and regulators of gene expression inside cells. RNAs undergo several metabolic events such as splicing, editing, localization, translation, and decay during their intracellular lifecycle. RNA metabolic events are executed by ribonucleoprotein complexes composed of RNA-binding proteins (RBPs), adapter proteins, and regulatory factors. Over 2,000 human genes encode proteins that are part of ribonucleoprotein complexes^1,2^. Individual RNA-associated proteins often regulate the metabolism of hundreds of RNAs. Mutations in RNA-associated proteins are associated with many human diseases including cancer, neurodegeneration, and developmental disorders^3,4^. Thus, decoding the effect of RBPs and associated factors on RNA metabolism is critical for our understanding of post-transcriptional gene regulation and molecular mechanisms underlying human disease.

Despite extensive biochemical studies of RNA metabolism and RBP function, we do not know the full set of cellular factors that regulate specific RNA metabolic events. This is because binding of RBPs can increase, decrease or leave unchanged metabolic events on their target RNA depending on their affinity, location, and other associated factors^5–8^. Many RBPs also associate with multiple ribonucleoprotein complexes and participate in several distinct RNA metabolic events^9^. Conversely, protein factors that do not directly bind RNA can still affect RNA metabolism by regulating the interactions between RNAs and RBPs, or by controlling the cellular level and activity of RBPs^10^. Hence, biochemical studies of RBP-RNA interactions are insufficient to reveal the full spectrum of functional regulators of RNA metabolic events in cells.

Unbiased genetic screening can identify cellular factors regulating RNA metabolism, but are limited in their current form. CRISPR screens using indirect phenotypes such as cell growth and fluorescent protein levels are difficult to engineer and interpret for many RNA metabolic events^11,12^ due to potential false positives^13,14^ and genetic compensatory mechanisms^15^. CRISPR perturbations followed by arrayed bulk RNA sequencing or pooled single cell RNA sequencing can directly report on RNA phenotypes^16,17^. But these transcriptome-wide sequencing approaches have limited flexibility to study different types of RNA metabolic events, are biased towards highly expressed RNAs, and are costly and labor intensive to scale beyond a few dozen perturbations. Thus, it has not been possible until now to genetically dissect the RNA-centric functions of human proteins at the scale of reporter-based CRISPR screens and with the ability to directly monitor diverse RNA metabolic events.

## Results

### Development of RNA-linked CRISPR screening in human cells

We reasoned that combining CRISPR-based perturbations with barcoded RNA readouts will provide a general approach to study the genetic control of different events in human RNA metabolism. Supporting the feasibility of this barcoding approach, RNA interference screens in human cells^18^ and CRISPR interference screens in *S. cerevisiae*^19,20^ have used barcoding to link perturbations to transcriptional readouts. However, lentiviral delivery, commonly used for CRISPR screening in human cells, will scramble sgRNA-barcode linkages due to template switching during reverse transcription^21–23^ and result in variable expression of RNA barcodes due to random genomic integration^24–26^. To avoid these limitations, we employed an iterative, site-specific integration strategy to stably express SpCas9 (Cas9 hereafter), sgRNAs, and barcoded RNA reporters from a defined genomic locus (Fig. 1A). First, we generated a clonal HEK293T cell line with a single *attP* ‘landing pad’ site for the Bxb1 integrase^27,28^ at the *AAVS1* safe harbor locus by Cas9-mediated homology-directed repair. Next, we integrated a doxycycline-inducible *Cas9* and an orthogonal *attP\** site^29,30^ into the landing pad using Bxb1-mediated recombination. Finally, we integrated sgRNA and reporter RNA cassettes into the *attP\** site using Bxb1-mediated recombination. We used fluorescent and antibiotic selection markers to enrich for cells with successful integration events at each step, and we used insulator elements to reduce transcriptional interference and promote long-term stable expression of integrated genes (Methods). Using an *EYFP* fluorescent reporter, we confirmed its uniform and stable expression after integration (Fig. 1B, blue). After doxycycline addition, we observed a progressive decrease in EYFP signal over 7 days that was specific to cells co-expressing an *EYFP*-targeting sgRNA (Fig. 1B, yellow), validating our ability to robustly induce Cas9-mediated gene knockout.

**Figure 1.**
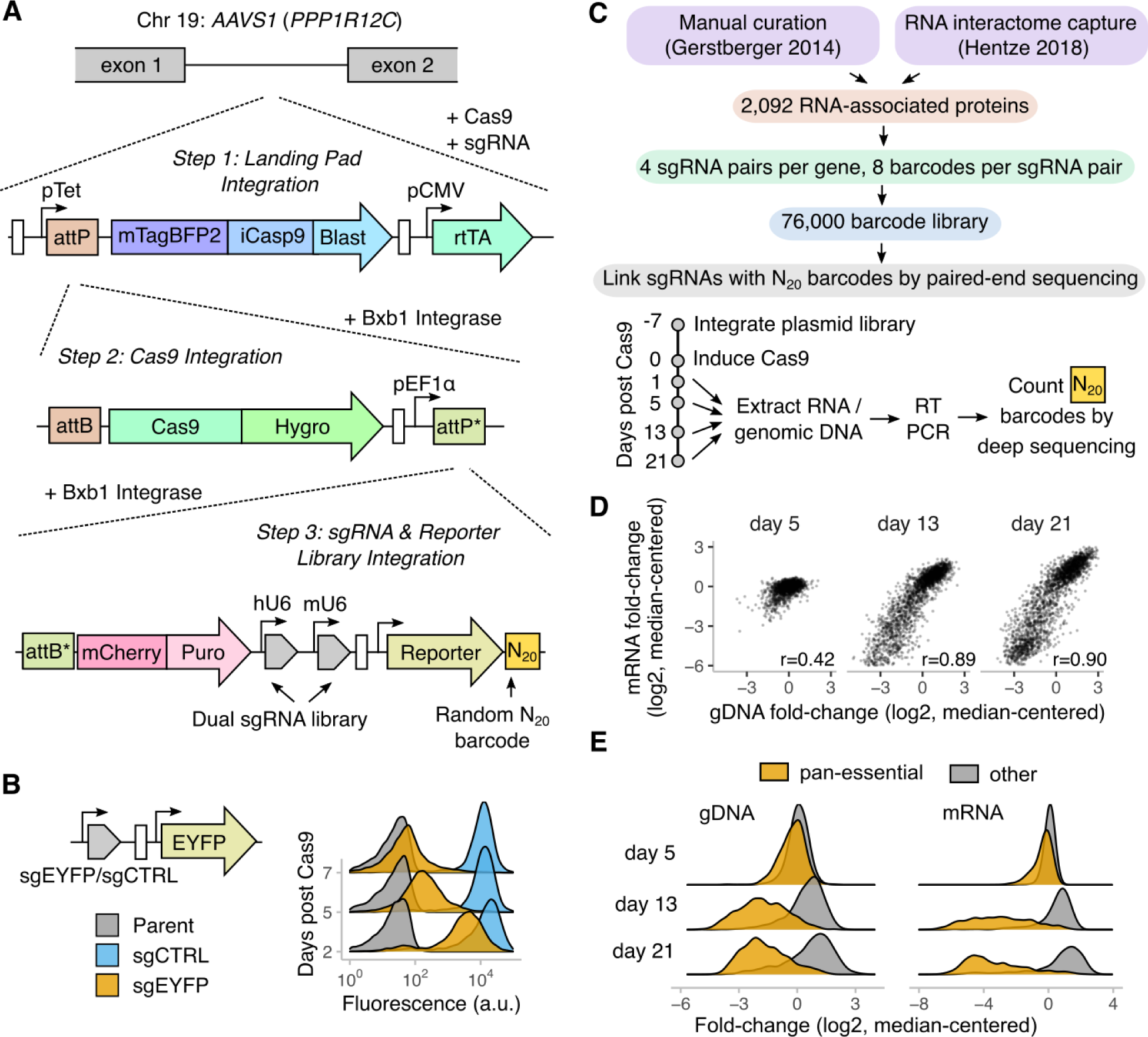
Development of RNA-linked CRISPR (ReLiC) screening in human cells. **A.** *Strategy for genomic integration of Bxb1 attP landing pad, SpCas9, and dual sgRNA and barcoded RNA reporters.* Unlabeled white rectangles represent cHS4 insulator sequences. *attP* and *attP\** refer to orthogonal recombination sites for the Bxb1 integrase that differ by a single nucleotide mismatch and undergo recombination only with their corresponding *attB* and *attB\** sites. Genetic elements are not drawn to scale. **B.** *Validation of Cas9 activity.* sgEYFP and sgCTRL are single guide RNAs targeting EYFP or a non-targeting control, respectively. Each histogram represents fluorescence of 10,000 cells as measured by flow cytometry. ‘Days post Cas9’ refers to days after addition of doxycycline to induce Cas9 expression. **C.** *Strategy for ReLiC sgRNA library design and validation.* sgRNAs and barcodes were iteratively cloned as shown in Fig. S1A and integrated into the genome as shown in Fig. 1A. **D.** *Correlated change in barcode frequency between genomic DNA and mRNA after Cas9 induction*. Each point corresponds to fold-change in mRNA or genomic DNA (gDNA) barcode counts for a single gene between day 1 and days 5, 13, or 21 post Cas9 induction. Fold-changes are median-centered across sgRNA pairs (sgRNAs henceforth) in the library, and the gene level fold-changes are median values across all detected sgRNAs for that gene. *r* refers to Pearson correlation coefficient between mRNA and genomic DNA log_2_ fold-changes. **E.** *Essential genes are depleted in genomic DNA and mRNA after Cas9 induction.* Histogram of fold-change in mRNA or genomic DNA counts for all genes targeted in the ReLiC library. Essential genes were defined as genes annotated as pan-essential in the DepMap database (n = 745). All other genes targeted in our library were classified as non-essential (n = 1401).

To identify regulators of RNA metabolism, we targeted 2,092 human genes encoding proteins annotated to interact with RNAs or RNA-binding proteins in previous manual curation and RNA interactome surveys^1,2^ (Fig. 1C). We selected sgRNAs from the validated Brunello library^31^ and used a dual sgRNA design to maximize knockout efficiency. We cloned the sgRNA pairs along with random N_20_ barcodes into a modular *attB\**-integrating vector that allows insertion of arbitrary RNA reporters (Fig. S1A). Our final library targeted 2,190 genes with 4 sgRNA pairs per gene, and included positive control sgRNA pairs targeting essential genes and non-targeting sgRNA pairs as negative controls (Table S1). We linked the N_20_ barcodes to sgRNAs by paired-end deep sequencing of the cloned plasmid library. We then integrated this library into our *attP\** parental cell line, and counted barcodes in the genomic DNA and transcribed RNA by deep sequencing (Fig. 1C, Fig. S1B). We recovered a median of 8 barcodes per sgRNA pair (henceforth referred to as sgRNAs) with at least one barcode for 99% of sgRNAs and 100% of all genes (Fig. S1C), thus capturing the diversity of our input library.

To test whether sgRNA-linked barcodes capture fitness effects, we sequenced and counted barcodes in genomic DNA and mRNA at different time points after Cas9 induction (Table S6). Barcode counts showed little systematic change on day 5 after Cas9 induction (Fig. 1D, left panel). However, on days 13 and 21 after Cas9 induction, barcode counts for a subset of sgRNAs were strongly depleted in both the genomic DNA and mRNA in a highly correlated manner (Fig. 1D, middle and right panels). The magnitude of depletion was correlated across distinct barcode sets for each gene (Fig. S1D), indicating the assay’s technical reproducibility. Barcodes in the genomic DNA corresponding to annotated essential genes (n = 745) were depleted 5.4–6.6 fold (median depletion at days 13 and 21) relative to other genes targeted in our library (n = 1401, Fig. 1E). Barcodes in the mRNA corresponding to the same essential genes were depleted 13.6–28.4 fold (median depletion at days 13 and 21) relative to other genes targeted in our library. The greater effect of essential gene knockout on mRNA relative to DNA might arise from the decreased RNA content in slower growing cells^32^. In summary, our RNA-linked screening strategy accurately captures both the identity and fitness effect of CRISPR perturbations solely from sequencing of barcodes in mRNA and genomic DNA.

### ReLiC identifies regulators of mRNA translation

We first applied ReLiC to study translation, an RNA metabolic step that is not directly accessible to existing CRISPR screening methods. The traditional gold standard for monitoring translation is polysome profiling — ultracentrifugation of cell lysate through a density gradient to separate mRNAs based on their ribosome occupancy^33–35^. We sought to combine this classic biochemical fractionation technique with ReLiC screening to identify regulators of ribosome occupancy on mRNAs. We used a spliced β-globin reporter^36,37^ as a prototypical model of a well-translated mRNA (Fig. 2A). First, to estimate the precision of our barcode-based measurement, we inserted 6 random barcodes into the 3′ UTR of this reporter and stably integrated the barcoded reporter pool into our *attP\** parental cell line. We counted the RNA barcodes in monosome (one ribosome), light polysome (2-4 ribosomes), and heavy polysome (>4 ribosomes) fractions by sequencing. Over 75% of the β-globin mRNA was in the light and heavy polysome fractions while the relative amount of the six barcodes varied less than 3% within each fraction (Fig. 2B). Next, we cloned the β-globin reporter into our ReLiC-RBP plasmid library, integrated the library into the *attP\** cell line, and induced Cas9 for 7 days before fractionating cell lysates (Fig. 2A). The duration of Cas9 induction was chosen to allow for sufficient protein depletion while preventing loss of essential gene knockouts. After counting sgRNA-linked barcodes in each fraction, we used MAGeCK^38^ to identify sgRNAs that significantly altered the ratio of barcodes between heavy (H) or light (L) polysomes and monosomes (M) (Table S7). To call a gene as a ‘hit’, we required that at least 3 sgRNAs for that gene had consistent positive or negative effects on polysome to monosome barcode ratios ^1^ and controlled the resulting false discovery rate (FDR) at 0.05 (Table S8).

**Figure 2.**
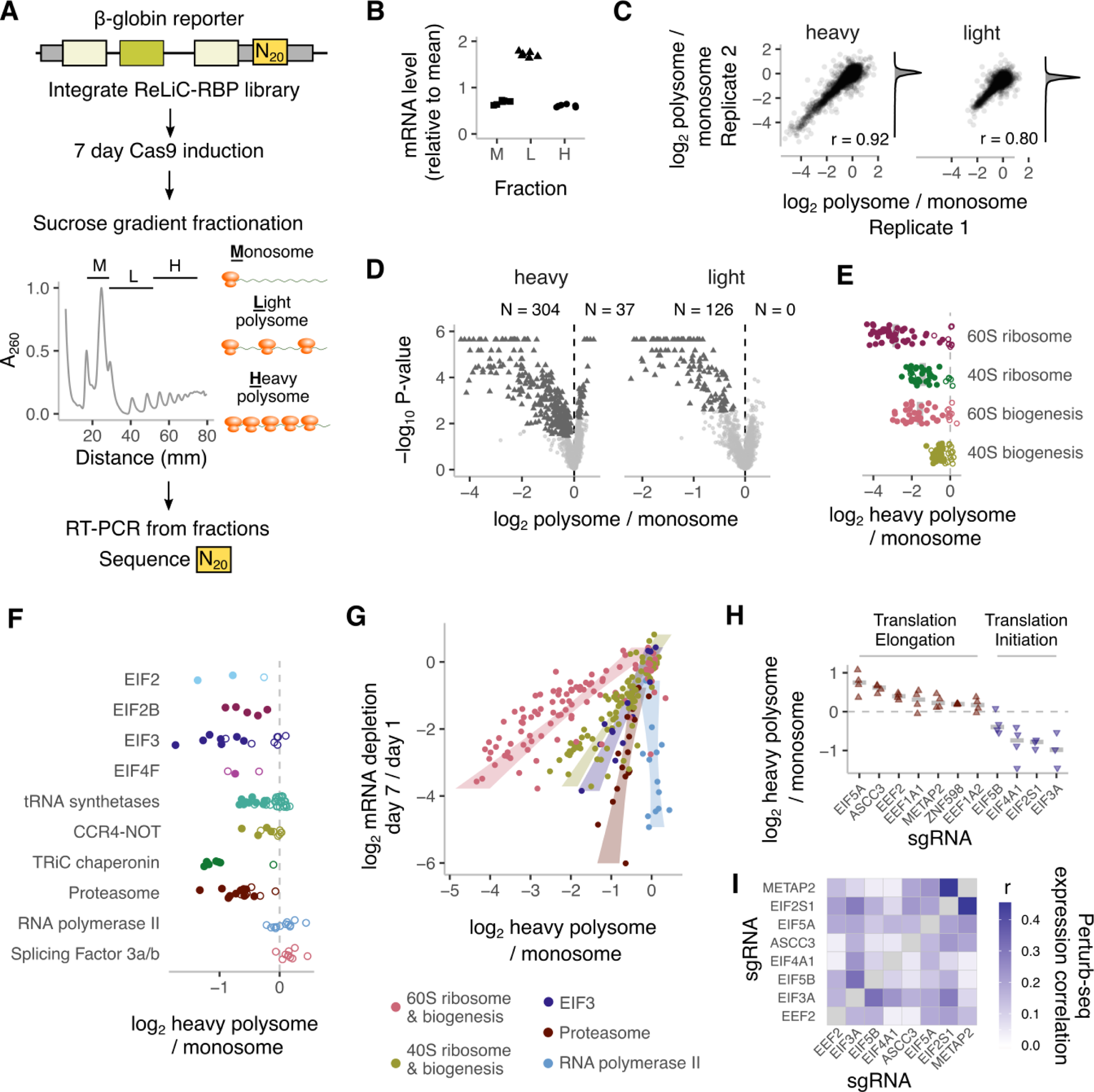
Polysome ReLiC identifies regulators of mRNA translation. **A.** *Strategy for combining ReLiC and polysome fractionation.* Lysates from cell pools expressing the ReLiC-RBP library with a β-globin reporter were fractionated on a 10-50% sucrose gradient to separate polysomes from monosomes. Absorbance at 260 nm (A_260_) was used to monitor ribosomal RNA signal along the gradient during fractionation. RNA extracted from monosome (M), light polysome (L), and heavy polysome (H) fractions was used to count reporter barcodes by deep sequencing. **B.** *Reporter distribution across polysome fractions.* Points correspond to relative mRNA level in each fraction for distinct 3′ UTR barcodes (n=6) for the β-globin reporter. **C.** *Correlation between replicates.* Points represent individual sgRNAs in the ReLiC library. Polysome to monosome ratios are median-centered across sgRNAs in the library. *r* refers to Pearson correlation coefficient. **D.** *Gene hits that alter polysome to monosome ratio.* Each point corresponds to a gene targeted by the ReLiC library. Horizontal axis indicates median of polysome to monosome ratios across all detected sgRNAs for each gene. Vertical axis indicates gene-level P-value from MAGeCK. Number of genes with FDR < 0.05 and decreased or increased polysome to monosome ratio are indicated with *N* and the individual genes are highlighted in dark grey triangles. All other genes are shown as light grey circles. **E.** *Change in polysome to monosome ratio for ribosomal protein and ribosome biogenesis genes.* Closed circles are genes that we call as gene hits (FDR < 0.05 with 3 or more concordant sgRNAs). Open circles are genes that do not pass our gene hit threshold. **F.** *Change in polysome to monosome ratio for protein groups and complexes.* Closed and open circles denote gene hits and non-hits similar to E. **G.** *Comparison of ribosome occupancy and mRNA depletion.* Points correspond to genes belonging to one of the highlighted groups. Shaded areas correspond to 95% confidence intervals for a linear fit of log_2_ polysome to monosome ratio to log_2_ mRNA depletion within each gene group. **H.** *mRNA ratios between polysome fractions for individual translation factors.* Each point corresponds to a distinct sgRNA pair for that gene. Grey bars denote median log_2_ ratio across all detected sgRNA pairs for that gene. **I.** *Correlation of expression profiles upon depletion of translation factors as measured by Perturb-seq.* Bulk expression profiles are from a previous genome-scale Perturb-seq (multiplexed perturbation and single cell RNA-seq) study^16^. *r* refers to Pearson correlation coefficient. EEF1A1, EEF1A2, and ZNF598 depletion did not have significant expression correlation with any of the other depletions, so they are excluded to visualize differences between the remaining factors.

Polysome to monosome ratios for individual sgRNAs were highly reproducible (r=0.92 and 0.80 for H/M and L/M, respectively) between replicate experiments (Fig. 2C). We identified 304 and 126 gene knockouts that decreased heavy polysome to monosome and light polysome to monosome ratios, respectively. 37 gene knockouts increased heavy polysome to monosome ratio, while none increased light polysome to monosome ratio (Fig. 2D). The skewed distribution of gene hits with more perturbations leading to a decrease in ribosome occupancy likely results from the efficient translation of β-globin mRNA in unperturbed cells (Fig. 2B). Consistent with heavy polysome fractions containing better-translated mRNAs, heavy polysome to monosome ratios were more sensitive to perturbations with more gene hits and larger effect sizes than light polysome to monosome ratios (Fig. 2D). We therefore focused on heavy polysome to monosome ratios for further analyses.

Gene hits that decreased polysome to monosome ratios were highly enriched for cytoplasmic ribosomal proteins and ribosome biogenesis factors (Fig. 2D, Fig. S2A). Indeed, 44 of the 54 large ribosomal (RPL) proteins and 28 of the 36 small ribosomal (RPS) proteins were classified as hits by MAGeCK (closed circles, Fig. 2E). As a group, knockout of large ribosomal proteins decreased polysome to monosome ratios more than knockout of small ribosomal proteins (Fig. 2E, median log_2_ H/M: −3.17 vs −1.48 for RPL vs RPS). Similarly, knockout of large ribosomal subunit biogenesis factors decreased polysome to monosome ratios more than knockout of small ribosomal subunit biogenesis factors (Fig. 2E, median log_2_ H/M −1.82 vs −0.55 for large vs small subunit biogenesis factors), though their overall effects were smaller than knockout of corresponding ribosomal proteins.

Translation initiation factors were also enriched among gene hits that decreased heavy polysome to monosome ratio (Fig. S2A), but their effects were more variable (Fig. 2F) and generally smaller than the effect of ribosomal protein depletion. Subunits of the EIF2, EIF2B, EIF3, and EIF4F initiation complexes all emerged as gene hits (closed circles, Fig. 2F). Some of the initiation factor subunits that we did not classify as hits (open circles, Fig. 2F) still had multiple sgRNAs that decreased heavy polysome to monosome ratio but fell just below our gene-level FDR threshold (EIF4G1 – FDR: 0.08) or did not meet our stringent criterion of 3 distinct sgRNAs with significant effects (EIF2S2, EIF4E – 2 sgRNAs). In the case of the 12-subunit EIF3 and associated EIF3J, the seven subunits A,B,C,D,E,G,and I that we called as hits were the same ones that severely reduced polysome to monosome ratio and fitness when depleted by siRNA in HeLa cells^39^. Aminoacyl-tRNA synthetase knockouts had mild and variable effects on ribosomal occupancy (Fig. 2F), presumably reflecting a balance between their direct effect on translation elongation and indirect effect on translation initiation through GCN2 and EIF2α phosphorylation^40–42^.

We identified several gene knockouts outside the core translation machinery with decreased polysome to monosome ratio (Fig. 2F). Four subunits of the CCR4-NOT complex (CNOT1, CNOT2, CNOT3, and CNOT7), which has been implicated in a wide range of RNA metabolic processes^43^, emerged as hits in our screen, which agrees with observations in *S.cerevisiae* strains lacking CNOT2 and CNOT3 homologs^44^. Knockout of several subunits of the proteasome and the TRiC chaperonin complex led to substantially reduced polysome to monosome ratios, comparable in magnitude to knock-out of core translation initiation factors (Fig. 2F). Notably, these complexes did not arise as hits simply because of their essentiality since knockout of other essential cellular complexes such as RNA polymerase II and splicing factor 3a/b (SF3) did not reduce polysome to monosome ratio (Fig. 2F). While neither the proteasome nor the TRiC chaperonin complex has been directly associated with translational regulation, they play a critical role in maintaining cellular proteostasis by coordinating their activities with translational output^45,46^. Our results reveal a reciprocal regulation of translation in response to changes in proteasomal and chaperonin capacity.

### Pathway- and mechanism-specific effects of gene knockouts on ribosome occupancy

Ribosome occupancy on mRNAs is often correlated with cellular growth rate, with slower growth accompanied by lower polysome to monosome ratio across different growth conditions and organisms^39,47,48^. Our measurements of both barcode depletion after Cas9 induction and polysome distribution of barcodes allowed testing the generality of the relationship between ribosome occupancy and growth across ∼2,000 different gene perturbations. Across all perturbations, decrease in polysome to monosome ratio was positively correlated with barcode depletion in both mRNA and genomic DNA but had a wide distribution (Fig. S2B). We then focused on gene knockouts for ribosomal proteins, ribosome biogenesis factors, EIF3 subunits, proteasome, and RNA polymerase since these groups have several essential genes with varying growth effects. Within each group, gene knockouts with lower polysome to monosome ratio also showed depleted mRNA and genomic DNA (Fig. 2G, Fig. S2C). However, each gene group had characteristically distinct relationship between ribosome occupancy and growth fitness as measured by barcode depletion. Perturbing large ribosomal proteins and biogenesis factors resulted in the largest decrease in polysome to monosome ratio relative to fitness, which was followed by perturbations of small ribosomal proteins and biogenesis factors, and then EIF3 (Fig. 2G, Fig. S2C). Perturbing proteasomal subunits produced a smaller but still significant decrease in ribosome occupancy, while perturbing RNA polymerase II subunits did not alter ribosome occupancy despite their significant effects on growth fitness (Fig. 2G, Fig. S2C). Hence, the coupling between growth rate and ribosome occupancy in human cells is not invariant across all perturbations, but depends on the pathway or the molecular process that is limiting growth.

We next examined the small group of gene knockouts that increased the heavy polysome to monosome ratio in our screen (Fig. 2H, brown triangles, Table S8). The translation factors EEF2 and EIF5A were among our top hits, consistent with their known role in promoting translation elongation. Knockout of the canonical elongation factors EEF1A1 and EEF1A2 also significantly (P = 0.03-0.04) increased ribosome occupancy though they fell just below our FDR threshold for calling gene hits (FDR = 0.08-0.09). Intriguingly, the ribosome-associated quality control factor ASCC3 was the top gene hit for increased heavy polysome to monosome ratio (log_2_H/M = 0.62, FDR = 1e-4). Since ASCC3 is involved in splitting stalled ribosomes on mRNAs^49^, its presence here suggests that even well-translated mRNAs such as this β-globin reporter undergo some degree of ribosome stalling and quality control. Supporting this inference, knockout of the ribosome collision sensor ZNF598, which acts upstream of ASCC3^49^, also increased ribosome occupancy (log_2_H/M = 0.19, FDR = 0.06, p = 0.007). In addition, knockout of METAP2, which removes methionine from the N-terminus of nascent polypeptides, increased ribosome occupancy (log_2_H/M = 0.22, FDR = 0.001, p = 3e-4), pointing to an effect of nascent peptide processing on the kinetics of mRNA translation.

Finally, we asked whether differential effects of gene perturbations on ribosome occupancy as measured by polysome to monosome ratios are reflected in their cellular transcriptional response. Using a genome-scale Perturb-seq dataset^16^, we correlated and clustered the transcriptional profiles of translation factor perturbations that had concordant or discordant effects on ribosome occupancy (Fig. 2I). Perturbations with concordant effects on ribosome occupancy (Fig. 2H) did not show a higher correlation with each other than with perturbations with discordant effects on ribosome occupancy. In fact, depletion of METAP2 and EIF2S1 (EIF2α), which are known to interact at a molecular level^50^, had a markedly higher correlation in their transcriptional response even though these gene knockouts had discordant effects on ribosome occupancy (Fig. 2H). Thus, the effects on ribosome occupancy measured by ReLiC are distinct from the downstream transcriptional responses to these perturbations.

### Isoform-selective splicing screens using ReLiC

We next applied ReLiC to investigate regulators of alternative splicing, an RNA processing event that occurs on most endogenous human mRNAs^53^. Existing screening approaches to study splicing require careful design of fluorescent protein reporters^13,54^ and can result in high false positive and negative rates^14^. We reasoned that insertion of barcodes in the 3′ UTR will allow us to directly measure the ratio of different splice isoforms carrying the barcode and thus capture the effect of the linked sgRNA perturbation in our ReLiC screens. To test this idea, we used the same β-globin reporter^36,37^ as in our translation screen (Fig. 3A). RNA-seq of cells stably expressing the β-globin reporter confirmed that the canonically spliced β-globin mRNA with three exons is by far the most abundant isoform with less than 1% of reads mapping to the two introns or to the splice junction between exons 1 and 3 (Fig. 3B). We then performed three isoform-selective screens for regulators that increase intron 1 retention (i12), intron 2 retention (i23), or exon 2 skipping (e13) (Fig. 3A). After harvesting RNA 1, 3, 5 and 7 days post Cas9 induction, we selectively amplified each isoform along with the barcode using primers that anneal to the two introns or to the exon 1-exon 3 junction (Fig. 3C) and counted barcodes by deep sequencing. Similar to our polysome ReLiC screen, we used an FDR threshold of 0.05 and a minimum of three concordant sgRNAs for calling gene hits.

**Figure 3.**
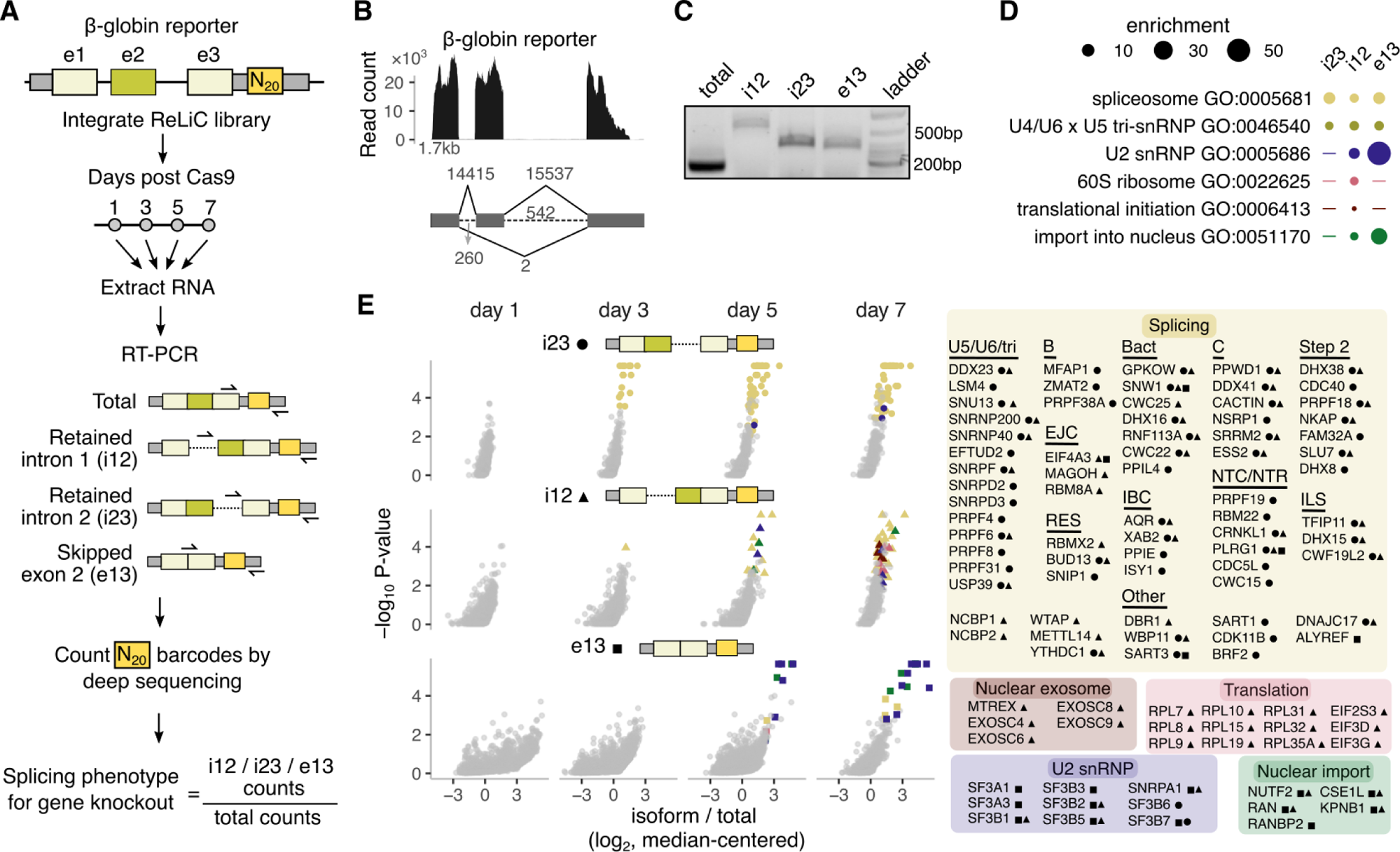
Isoform-specific splicing screens using ReLiC. **A.** *Schematic of ReLiC splicing screens.* A β-globin reporter with 3 exons (e1, e2, e3) separated by two introns was used for ReLiC splicing screens. After ReLiC library integration and Cas9 induction, RNA was harvested at different time points. Barcodes linked to different isoforms corresponding to retained intron 1 (i12), retained intron 2 (i23), skipped exon 2 (e13), or all isoforms (total) were amplified by PCR and counted by deep sequencing. Location of RT primer and PCR primers used for PCR amplification of barcodes for each isoform are shown as black arrows. Splicing phenotype for each gene was calculated as the log_2_ ratio of barcode counts for each isoform to the total barcode counts using MAGeCK. Isoform ratios are median values across all sgRNAs for each gene after median-centering across all sgRNAs in the library. **B.** *Relative abundance of reporter splice isoforms.* Top panel shows RNA-seq read count at each nucleotide position of the 1.7kb β-globin reporter. Bottom panel shows the different splice isoforms and the read counts mapping to each splice junction or intron. **C.** *Selective amplification of barcodes linked to splice isoforms.* Agarose gel lanes show RT-PCR products of expected size for the different isoforms: total: 181bp, i12: 532bp, i23: 302bp, and e13: 286bp. **D.** *Gene ontology analysis*. Selected cellular processes and components enriched among gene hits on day 7 after Cas9 induction. Markers are sized according to the fold enrichment of the GO term. GO terms with FDR > 0.05 are indicated by dashes. **E.** *Identity of gene hits.* Each point corresponds to a gene targeted by the ReLiC library. Different panels correspond to days after Cas9 induction (horizontal) and isoform screens (vertical). Marker shape denotes isoform identity and marker color denotes one of five highlighted gene groups. Genes with FDR < 0.05 and belonging to one the highlighted groups are listed in the legend.

We detected few or no gene hits one day after Cas9 induction for the three splice isoforms (Fig. S3A, Fig. 3E), consistent with few proteins being depleted at this early time point after their gene knockout. As the duration of Cas9 induction increased, the three isoforms exhibited markedly distinct responses (Fig. S3A, Fig. 3E). Three days after Cas9 induction, 18 gene knockouts increased intron 2 (i23) retention while few or no gene hits were detected for the intron 1-retained and the exon 2-skipped isoforms. This difference between isoform levels persisted at day 5, suggesting that splicing of intron 2 is more sensitive to genetic perturbations than the other two isoforms. A larger number of gene hits emerged for the intron 1-retained isoform by day 7 (N = 101), while the number of gene hits for the intron 2-retained isoform remained similar between days 5 and 7 (N = 62 and 54). Fewer gene knockouts increased the exon 2-skipped isoform (N = 22, 25 at days 5, 7) in comparison to the two intron-retained isoforms at all time points. Effect sizes of the gene hits were reproducible across distinct barcode sets for each gene (Fig. S3B) and specific to each isoform (Fig. S3C).

The three isoform-specific screens identified both common and unique sets of gene hits that were evident by automated gene ontology analysis (Fig. 3D) and by manual inspection (Fig. 3E). Gene hits in the two intron retention screens were dominated by core spliceosome components and splicing-associated factors (yellow circles and triangles, Fig. 3E). Spliceosome hits were distributed throughout the splicing cycle starting from the tri-snRNP complex that is required to form the catalytically active spliceosome and included members from most known spliceosomal subcomplexes^55–58^. Our screen also identified *trans* regulators of spliceosomal function such as CDK11B – a recently identified activator of the SF3b complex^59^, and BRF2 – an RNA polymerase III subunit required for transcription of U6 snRNA^60^.

Retention of intron 1 was promoted by an additional group of gene knockouts that were enriched for mRNA translation and nuclear RNA exosome factors (red and brown triangles, Fig. 3D,E). Loss of ribosomal proteins and translation factors might inhibit nonsense-mediated decay (NMD) of the intron 1-retained isoform. While retention of either intron 1 or intron 2 will generate a premature termination codon (PTC), only the intron 1-retained isoform will have a splice junction and an associated exon-junction complex (EJC) downstream of the PTC, which is a well-known trigger for NMD^61–64^. Consistent with a role for NMD, EJC components (MAGOH, EIF4A3, RBM8A) and RNA export factors (NCBP1, NCBP2) also emerged as hits only in the intron 1 retention screen (Fig. 3E). Nevertheless, core NMD factors such as UPF and SMG proteins were not detected in any of the splicing screens, while the effect of nuclear RNA exosome components might be indirect through their role in ribosome biogenesis or RNA export^65,66^.

### Differential effects of SF3b complex subunits on splicing

In contrast to intron retention, perturbations increasing exon 2 skipping were enriched for a narrow set of splicing factors. Components of the U2 snRNP, most notably several members of the SF3 complex, were among the top hits (purple squares, Fig. 3D,E), suggesting that their depletion allows some degree of splicing but impairs the correct selection of splice sites. This is consistent with the subtle alterations in exon skipping caused by disease-causing mutations in the SF3b complex^67–69^. Exon 2 skipping was also promoted by perturbing components involved in nuclear protein import (green squares, Fig. 3D,E), presumably through their effect on nuclear import of U2 snRNP proteins after their synthesis in the cytoplasm. Perturbing individual components of the 7-subunit SF3b complex^70^ had distinct effects on exon skipping and intron retention (Fig. 4A), even though all 7 subunits are essential for cell growth (Fig. S3D). Exon 2 skipping was greatly increased upon loss of the subunits SF3B1, SF3B2, SF3B3, SF3B5, slightly increased by loss of SF3B7, and unaffected by loss of SF3B4 and SF3B6 (Fig. 4A). Intron 2 retention was increased by loss of SF3B6 and SF3B7, while intron 1 retention was increased by loss of SF3B1, SF3B2, and SF3B5 (Fig. 4A). By contrast, loss of the activating helicase AQR increased the retention of both introns 1 and 2 (brown markers, Fig. 4A).

**Figure 4.**
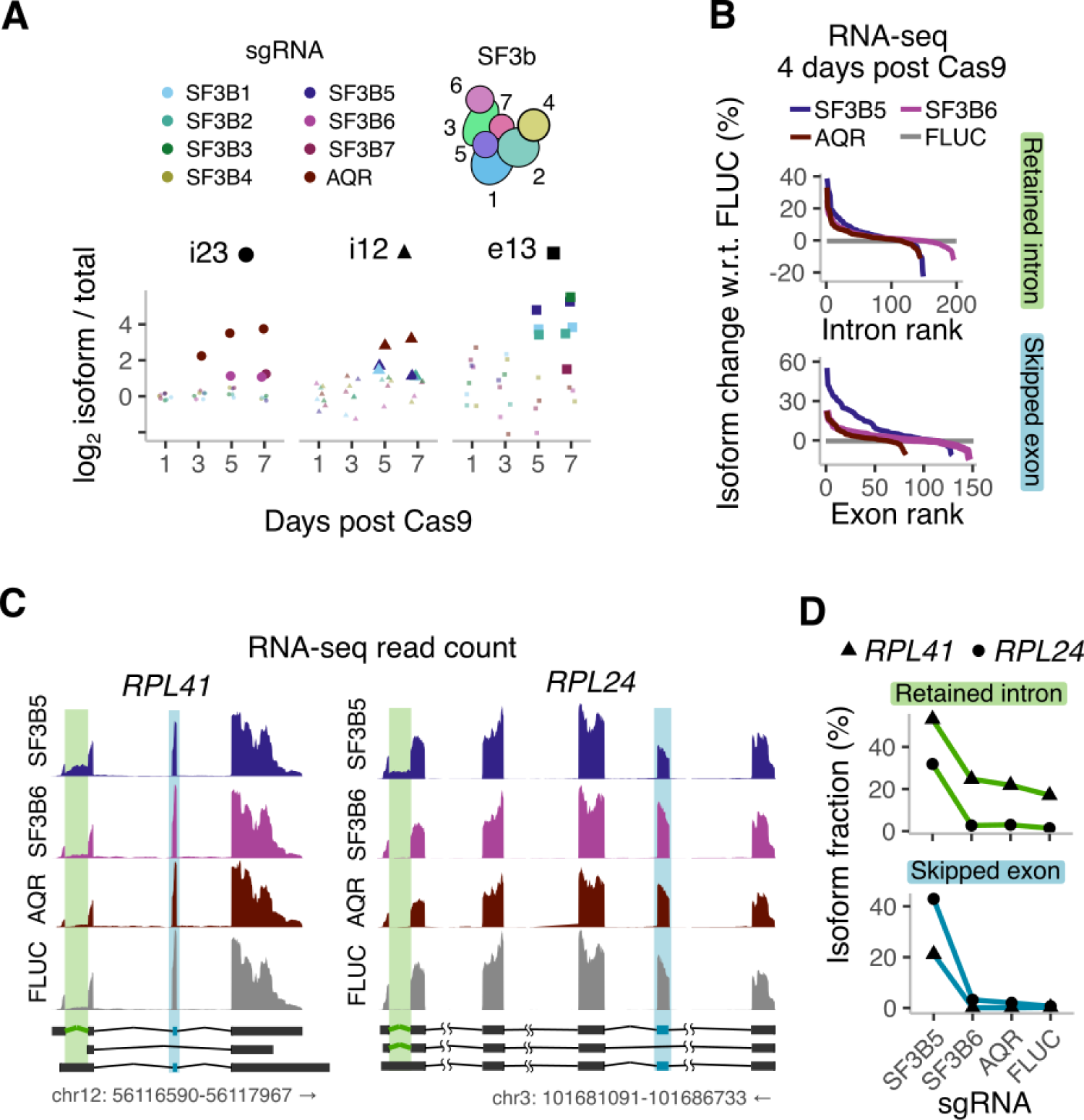
Differential effects of SF3b complex subunits on splicing. **A.** *Relative reporter isoform levels upon SF3b complex perturbations.* Splicing phenotypes are shown for genes encoding SF3b complex subunits and the helicase AQR. AQR is shown as a positive control hit for intron retention. FDR < 0.05 is indicated by large marker, and FDR ≥ 0.05 is indicated by small marker. **B.** *Change in endogenous splicing isoforms upon SF3b complex perturbations.* RNA-seq was performed 4 days after inducing Cas9 in cells expressing sgRNAs targeting SF3B5, SF3B6, AQR, or a non-targeting FLUC control. Change in intron retention or cassette exon skipping were calculated across all ENSEMBL-annotated transcripts, and ranked by decreasing magnitude of change with respect to the FLUC control sample. **C.** *Examples of endogenous isoform changes.* Read counts for *RPL41* and *RPL24* loci are shown for the RNA-seq from B. Specific retained introns and skipped exons are highlighted in green and blue rectangles, respectively. Schematics at the bottom correspond to ENSEMBL isoforms with the highlighted retained intron and skipped exon events. **D.** *Quantification of isoform fraction* for the endogenous intron retention and exon skipping events in C. Note that the RNA-seq coverage at the skipped exon in C reflects the magnitude of exon inclusion and not the magnitude of exon skipping.

We next examined how the differential effects of SF3b subunit depletion on β-globin reporter splicing extend to endogenous mRNAs. To this end, we generated HEK293T cell lines with the subunits SF3B5 and SF3B6, which affected distinct splicing events in our screen, individually depleted through Cas9-mediated knockout. We also targeted AQR, a top hit in both our intron retention screens, as a positive control and included a non-targeting control sgRNA against firefly luciferase (FLUC). We performed RNA-seq 4 days after Cas9 induction to identify endogenous splicing events that are particularly sensitive to the respective genetic perturbations. Loss of SF3B5 increased skipping of 45 annotated cassette exons by 10% or higher (Fig. 4B). For some cassette exons, the exon skipped isoform increased over 10-fold from less than 2% to 20-40% of the total isoform fraction (Fig. 4C,D). Loss of SF3B6 or AQR affected the skipping of less than 10 cassette exons at the same effect size, while all three splicing factors increased aberrant retention of a similar number of distinct introns (Fig. 4B, Fig. S3D). Interestingly, increased intron retention and exon skipping upon SF3B5 loss occurred at distinct splice sites within the same transcriptional unit for genes such as RPL24 and RPL41 (Fig. 4D). In summary, the differential effects of SF3b subunits on splicing of the β-globin reporter extend to endogenous mRNAs with a subset of SF3b subunits playing a more prominent role in regulating exon skipping.

### ReLiC screen for regulators of mRNA quality control

Our finding of ribosomal proteins and core translation factors as hits in our screen for intron retention (Fig. 3D,E) suggest that they promote the decay of aberrantly spliced mRNAs through the NMD pathway. However, previous CRISPR screens for NMD using fluorescent protein reporters recovered few ribosomal proteins and core translation factors^71,72^, presumably because these genes are critical for protein expression. We reasoned that sequencing mRNA barcodes using ReLiC provides a general approach to identify regulators of mRNA quality control pathways independent of their effect on protein expression. To test this idea, we modified the β-globin reporter from previous screens to add a premature termination codon (PTC) at position 39 in the second exon (Fig. 5A), such that it is similar to previously used reporters for NMD^36^. At steady state, mRNA levels of the PTC-containing reporter were strongly reduced relative to a reporter with a normal termination codon (NTC, Fig. S4A). To measure mRNA effects specific to the PTC and NTC reporters, we combined our ReLiC-RBP library with a dual barcoding strategy^19^ to normalize barcode counts for the reporter of interest relative to that of the mCherry-puro selection marker within each cell (Fig. 5A). We harvested RNA 7 days after Cas9 induction and counted mRNA barcodes for the PTC and NTC β-globin reporters and the mCherry-puro marker by deep sequencing.

**Figure 5.**
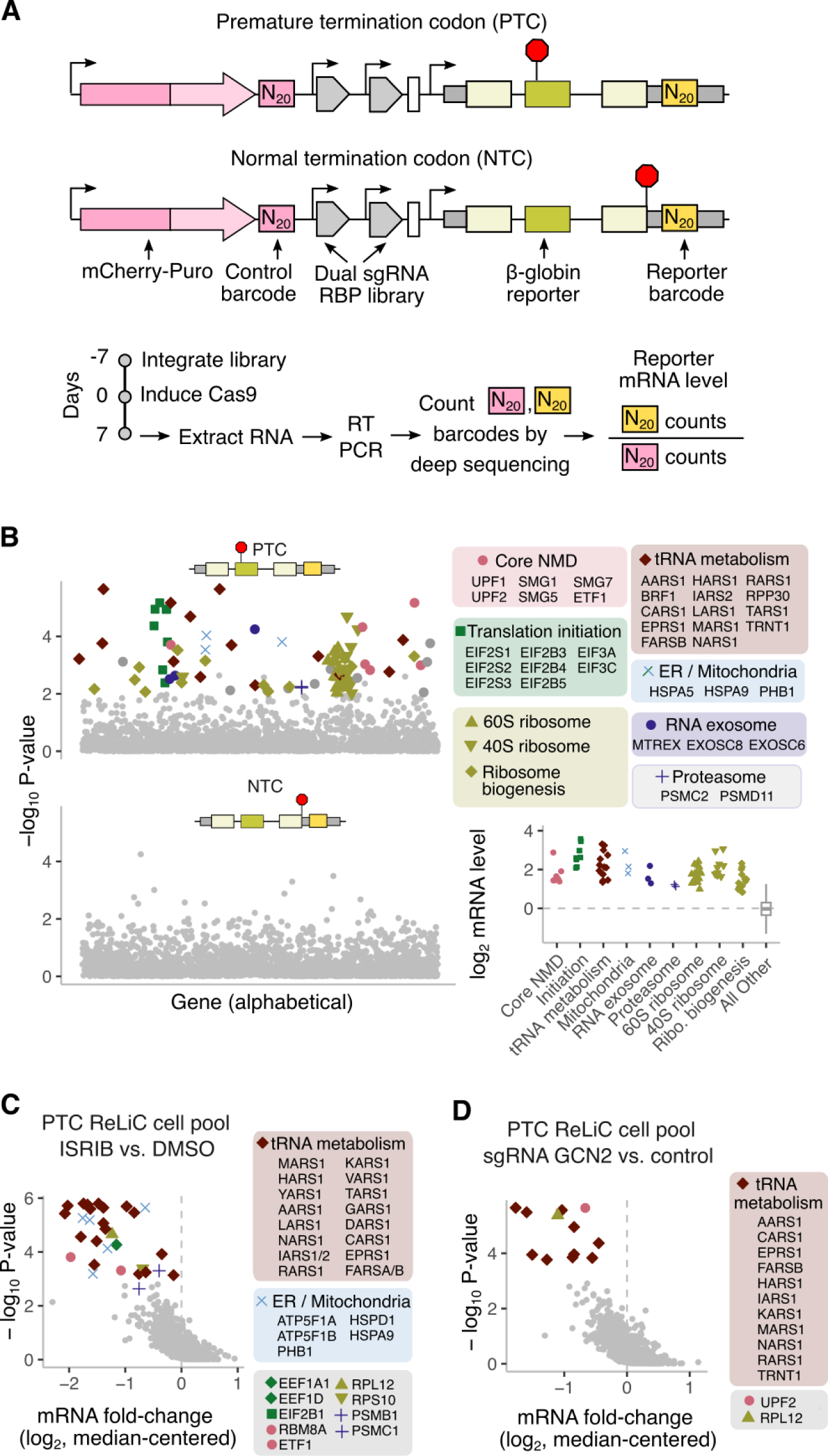
Dissecting co-translational quality control using chemogenomic ReLiC screening. **A.** *Dual barcode strategy for measuring reporter mRNA levels.* Red octagons represent location of stop codons along the β-globin reporter. N_20_ barcodes are added to the 3′ UTR of both the reporter and the mCherry-puro control. Reporter mRNA levels are calculated as the ratio of barcode counts for the reporter to the mCherry-puro control. Reporter mRNA levels represent median values across all sgRNAs for each gene and are median-centered across all sgRNAs in the library after log_2_ transformation. **B.** *Gene hits from dual barcode NMD screen.* Genes with increased reporter mRNA level are classified as hits if they have FDR < 0.05 as calculated by MAGeCK. Hits within one of the six highlighted gene groups are listed in the legend. Genes are arranged alphabetically along the x-axis. The lower right panel shows reporter mRNA level of the highlighted hits for the PTC reporter. Markers for gene hits are jittered along the x-axis to reduce overlap. **C.** *Chemical modifier screen with ISRIB.* Cell pool expressing the ReLiC-RBP library with the PTC reporter from a was treated with 200 nM ISRIB or DMSO for 48 hours after Cas9 induction for 5 days. mRNA fold-change is calculated by normalizing the barcode counts for each sgRNA in the ISRIB-treated sample to the corresponding counts in the DMSO-treated sample, and median-centered across all sgRNAs. Genes with lower mRNA level in the ISRIB-treated sample and FDR < 0.01 as calculated by MAGeCK are classified as hits. Marker colors and shapes denote the highlighted gene groups from B. **D.** *Genetic modifier screen with GCN2 depletion.* Cell pool expressing the ReLiC-RBP library with the PTC reporter from a was transduced with lentivirus expressing a GCN2-targeting sgRNA or a control sgRNA, followed by Cas9 induction for 7 days. mRNA fold-change is calculated by normalizing the barcode counts for the GCN2 sgRNA sample to the corresponding counts in the control sgRNA sample, and median-centered across all sgRNAs. Genes with lower mRNA level in the GCN2 sgRNA sample and FDR < 0.01 as calculated by MAGeCK are classified as hits. Marker colors and shapes denote the highlighted gene groups from B.

Our dual barcode ReLiC screen recovered 90 gene hits (FDR < 0.05, 3 sgRNAs with concordant effects) whose knockout increased levels of the PTC reporter relative to the mCherry-puro marker (Fig. 5B, Fig. S4C). We did not observe any hits for the NTC reporter at the same FDR threshold, as we would expect given that both the NTC reporter and mCherry-puro marker encode mRNAs with normal stability (Fig. 5B, Fig. S4C). Several core components of the NMD pathway (UPF1, UPF2, SMG1, SMG5, SMG7, ETF1) were among the gene hits for the PTC reporter, indicating our ability to identify NMD-specific factors (Fig. 5B, pink circles). Other NMD-associated factors such as SMG6 and EIF4A3 fell just below the FDR threshold but still significantly (MAGeCK P-value < 0.05) increased mRNA levels of the PTC reporter (Table S8). Remarkably, a large proportion of gene hits for the PTC reporter encoded core factors involved in various steps of mRNA translation (Fig. 5B, squares, triangles, and diamonds; Fig. S4B). These included both small and large ribosomal proteins, ribosome biogenesis factors, translation initiation factors, and aminoacyl-tRNA synthetases. These translation-related hits are consistent with the known requirement of mRNA translation for NMD^73^. Interestingly, several translation initiation factors in the EIF2, EIF2B, and EIF3 complexes emerged as hits in our NMD screen, while gene knockouts encoding the EIF4F complex (EIF4A1, EIF4E, EIF4G1) did not increase PTC reporter levels (Fig. S4D). Notably, the lack of EIF4F hits in our NMD screen was not simply due to variable knockout efficiency since EIF4F components had a similar growth depletion upon knockout as several EIF2, EIF2B, and EIF3 components (Fig. S4E). While the biochemical requirement for EIF4F in NMD remains unclear^74–76^, our genetic screen results suggest a limited *in vivo* role for EIF4F compared to EIF2, EIF2B, and EIF3 in regulating NMD.

### Chemical and genetic modifier screens using ReLiC

Our NMD screen also identified gene hits involved in ER and mitochondrial homeostasis (Fig. 5B, x markers). Since disruption of ER and mitochondrial homeostasis are known to trigger phosphorylation of EIF2α by the kinases PERK and HRI, our ER- and mitochondria-related hits might arise from phosphorylation of EIF2α upon their depletion. This is consistent with the known inhibition of NMD caused by phosphorylation of EIF2α^77–79^. To directly identify regulators of NMD that act through EIF2α phosphorylation, we adapted ReLiC to perform a chemical modifier screen using the small molecule ISRIB that renders translation insensitive to EIF2α phosphorylation^80^. After inducing Cas9 for 6 days, we treated a ReLiC cell pool expressing the PTC reporter with ISRIB or DMSO for 48 hours then harvested RNA and counted barcodes. We identified 30 gene knockouts (FDR < 0.01) that decreased mRNA levels of the PTC reporter upon ISRIB treatment relative to the DMSO control (Fig. 5C). These gene hits included several ER- and mitochondrially-localized proteins (Fig. 5C, x markers), consistent with their knockout inhibiting NMD through EIF2α phosphorylation. Some of the ISRIB screen hits were not identified in the original NMD screen since they fell just below the FDR threshold (Table S8).

Knockout of several aminoacyl-tRNA synthetases also decreased PTC reporter levels upon ISRIB treatment (Fig. 5C, diamonds), suggesting that their depletion inhibits NMD through phosphorylation of EIF2α rather than by decreasing translation elongation. To test this hypothesis, we performed a genetic modifier screen using ReLiC to deplete the EIF2α kinase GCN2, which is activated by uncharged tRNAs that accumulate upon inhibition of aminoacyl-tRNA synthetases^40,41^. We transduced the ReLiC cell pool with lentivirus expressing sgRNAs targeting GCN2 or a non-targeting control, induced Cas9 for 7 days, then harvested RNA and counted barcodes. We identified 12 gene hits (FDR < 0.01) that decreased PTC reporter levels upon GCN2 depletion (Fig. 5D), out of which 10 were aminoacyl-tRNA synthetases (Fig. 5D, diamonds), confirming their action through GCN2-mediated EIF2α phosphorylation. Together, the above experiments show that chemical and genetic modifier screening using ReLiC can dissect the pathways through which gene knockouts affect RNA metabolic processes.

### GCN1 regulates cellular responses to the anti-leukemic drug homoharringtonine

Homoharringtonine (HHT) is an FDA-approved chemotherapeutic that targets the ribosome and is used to treat chronic and acute myeloid leukemias^81^. HHT binds to the large ribosomal subunit to arrest initiating ribosomes at start codons and inhibit protein synthesis^82,83^, but how cells respond to this translational arrest is not well understood. Given ReLiC’s ability to identify regulators downstream of both mRNA translation and chemical perturbations, we sought to use this approach to probe the cellular response to HHT treatment. To this end, we performed ReLiC-RBP screens using a simple reporter encoding EYFP (Fig. 6A). After inducing Cas9 for 7 days, we treated the cell pool with 1 μM HHT or DMSO for 6 hours before harvesting RNA and counting reporter barcodes.

**Figure 6.**
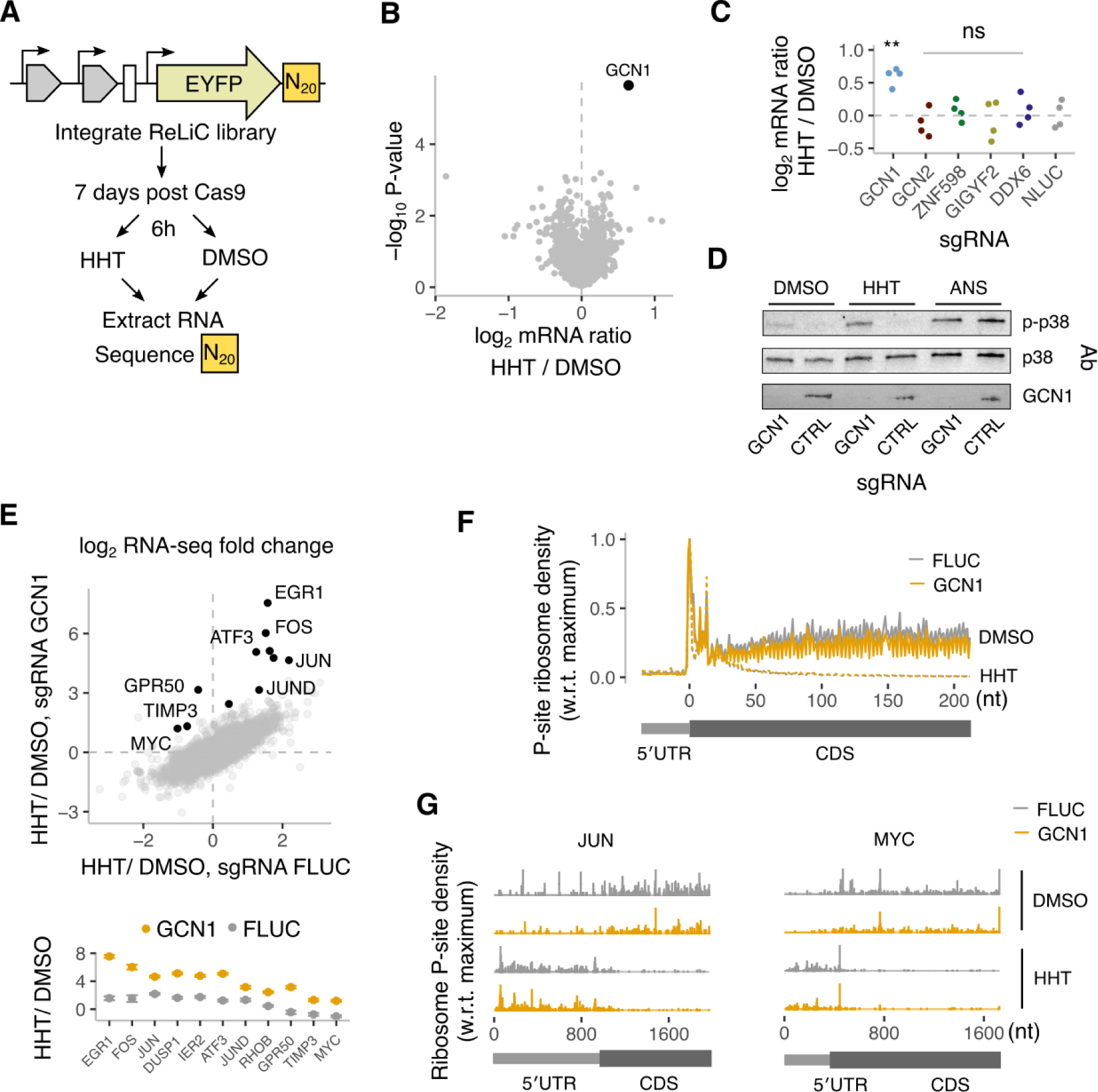
GCN1 regulates cellular responses to the anti-leukemic drug homoharringtonine. **A.** *Chemogenomic ReLiC screen using homoharringtonine (HHT).* ReLiC-RBP cell pool with an EYFP reporter was treated with 1 μM HHT or DMSO for 6 hours after Cas9 induction for 7 days. **B** *GCN1 regulates mRNA levels upon HHT treatment.* Each point represents a gene in the ReLiC-RBP library. Ratio of mRNA barcode counts for the reporter dre calculated between the HHT treatment and the DMSO-treated control, and are median-centered across all sgRNAs. Genes with increased reporter mRNA ratio are classified as hits if they have FDR < 0.05 as calculated by MAGeCK. **C.** *mRNA level changes upon HHT treatment for factors known to resolve ribosome collisions.* Points represent ratios between reporter barcode counts during HHT treatment compared to DMSO treatment for individual sgRNAs targeting each gene. P-values comparing the indicated perturbations to cells expressing the nontargeting Nluc control sgRNA are from a two sample t-test: ** (0.001 < P < 0.01), ns (P > 0.05). **D.** *Immunoblots for phosphorylation of p38 in HEK293T cells +/− GCN1.* Cells were treated with homoharringtonine (1 μM), anisomycin (10 μM), or DMSO for 1 hour. Anisomycin (ANS) is a positive control for ribosome collision-induced p38 phosphorylation. **E.** *GCN1-dependent changes in endogenous mRNA expression during HHT treatment.* RNA-seq was performed 8 days after inducing Cas9 in cells expressing dual sgRNAs targeting GCN1 or a non-targeting FLUC control. Prior to harvest, cells were treated with HHT (1 μM) or DMSO for 6 hours. Each points corresponds to a gene and represents the ratio of mRNA levels between HHT and DMSO treatment. Black highlighted points correspond to immediate early genes (IEGs), which are also shown separately in the lower panel. **F.** *Metagene alignment of ribosome P-site density in 5′ UTR and CDS region across all transcripts.* Ribosome profiling was performed on +/− GCN1 cells after harringtonine (1μM) or DMSO treatment for 1 hour. **G.** *Ribosome P-site density in 5′ UTR and CDS region of JUN and MYC transcripts.* X-axis indicates position along the transcript in nucleotides.

Unlike our previous ReLiC screens where we uncovered multiple gene hits and RNA metabolic pathways, a single gene, *GCN1*, emerged as a clear hit (FDR < 0.05) whose knockout increased EYFP reporter mRNA levels during HHT treatment (Fig. 6B). GCN1 activates the kinase GCN2 to trigger EIF2α phosphorylation in response to amino acid limitation^84^. GCN1 also binds collided ribosomes on mRNAs^85–87^, which can trigger both degradation of the nascent peptide and the mRNA^88,89^. However, since HHT arrests ribosomes at the start codon, we would not expect amino acid limitation or ribosome collisions to occur under these conditions. Indeed, our ReLiC screen during HHT treatment did not identify the uncharged tRNA sensor GCN2 or the ribosome collision sensor ZNF598 and its downstream effectors GIGYF2 and DDX6 as hits (Fig. 6C). Since ribosome collisions also trigger the ribotoxic stress response through the kinase ZAKα that was not included in our original screen^85^, we measured p38 phosphorylation in wild-type and GCN1-depleted cells. HHT treatment increased p38 phosphorylation in GCN1-depleted cells while wild-type cells did not show a corresponding increase. By contrast, treatment with the elongation inhibitor anisomycin potently triggered p38 phosophorylation in both wild-type and GCN1-depleted cells (Fig. 6D).

Ribosome collisions induced by elongation inhibitors trigger upregulation of immediate early genes at the mRNA level^90^. To test if GCN1 regulates a similar gene expression program during HHT treatment, we performed RNA-seq on wild-type and GCN1-depleted cells after 6 hours of HHT treatment and compared to control conditions. HHT treatment caused widespread changes in mRNA levels in both wild-type and GCN1-depleted cells with ∼225 up-regulated genes and ∼450 down-regulated genes (> 2-fold change, p < 0.05). However, a small group of 60 genes, which included the immediate early genes, showed differential up-regulation in the GCN1-depleted cells in comparison to wild-type cells (Fig. 6E). These included genes such as *FOS*, *JUN*, and *ATF3*, which were 2-4 fold up-regulated in wild-type cells upon HHT treatment but were up-regulated 25-50 fold in GCN1-depleted cells. Other genes such as *MYC* and *TIMP3* that were mildly down-regulated in wild-type cells upon HHT treatment were instead 2-fold or more up-regulated in the GCN1-depleted cells. The transcriptional upregulation of immediate early genes along with increased p38 signaling in GCN1-depleted cells point to a potential role for GCN1 in mitigating ribosome collisions during HHT treatment.

To test for occurrence of ribosome collisions on endogenous mRNAs during HHT treatment, we first performed polysome fractionation from both wild-type and GCN1-depleted cells after 1 hour of HHT treatment (Fig S4F). Polysomes collapsed into monosomes upon HHT treatment, and the disome peak was of comparable intensity and nuclease sensitivity in both wild-type and GCN1-depleted cells. Additionally, ribosome profiling after 1 hour of HHT treatment showed no significant differences in average ribosome occupancy on mRNAs between wild-type and GCN1-depleted cells (Fig. 6F). Thus, ribosome collisions do not occur during HHT treatment at a scale that is detectable by bulk biochemical fractionation and do not alter global ribosome occupancy on mRNAs. Nevertheless, highly expressed immediate early genes such as *JUN* and *MYC* exhibited extensive ribosome density throughout their the 5′ UTR during HHT treatment (Fig. 6G), which was also recapitulated by analysis of previous ribosome profiling studies (Fig S4G). Furthermore, ribosomes initiate at multiple in-frame start codons even in the absence of HHT on mRNAs of several immediate early genes such as *JUN*, *MYC*, and *JUND*^91–93^. Together, these observations suggest that collisions occur on these mRNAs between upstream initiated ribosomes that have transitioned to elongation and HHT-arrested initiating ribosomes at downstream start codons, which are then sensed by GCN1.

## Discussion

In this study, we demonstrate ReLiC, an RNA-linked CRISPR screening platform for genetic dissection of diverse RNA metabolic processes in human cells. ReLiC enables measuring the effect of thousands of gene perturbations on mRNA translation, splicing, and decay – molecular processes that are not readily accessible to existing CRISPR screening methodologies. Our work reveals networks of molecular pathways, protein complexes, and individual proteins that mediate the effect of *cis* sequence elements and chemical perturbations on RNA metabolism. The resulting effects are consistent with known molecular mechanisms and also provide new insights into the interplay between RNA metabolic processes and cellular physiology.

Combining ReLiC with biochemical fractionation reveals characteristic relationships between mRNA translation and other cellular processes. Knocking out proteasomal subunits decreases ribosome occupancy at a distinct rate relative to growth fitness. The robustness of this relationship hints at a rheostat that tunes the rate of global protein synthesis to match proteasomal capacity, and could be mediated by shared cellular signaling or metabolic pathways^94,95^. Conversely, the lack of effect of RNA polymerase II depletion on ribosome occupancy points to a tightly coordinated synthesis of the entire translation machinery at different rates of transcription. This decoupling between transcriptional capacity and ribosomal activity might enable human cells to maintain optimal rates of protein synthesis across diverse cell states and growth conditions, akin to bacteria^47,96^.

ReLiC reveals the role of essential pathways and genes in RNA metabolism even when their knockout is deleterious to cell growth. Chemical perturbations that abrogate protein expression can still be probed for their genetic dependencies using ReLiC, as demonstrated by our identification of GCN1’s role during HHT treatment. ReLiC captures differential effects of perturbations within the same protein complex such as between members of the SF3b complex and between large and small ribosomal proteins. Unlike biochemical strategies, ReLiC identifies both direct effectors and indirect regulators of RNA metabolism, as exemplified by the identification of translation-related pathways across our screens for ribosomal occupancy, splicing, and mRNA decay. In contrast to single cell screening approaches, ReLiC can straightforwardly combine CRISPR screening with bulk biochemical readouts of RNA metabolism, thus providing a powerful framework to access and screen for RNA phenotypes such as localization^97^, condensation^98^, and editing^99^. Further, ReLiC’s ability to selectively amplify and dissect rare RNA splicing events underscores its exquisite sensitivity and large dynamic range.

We anticipate that ReLiC can be extended to a broad range of biological settings, genetic perturbations, and RNA types. Applying ReLiC to diverse cell types, cell states, and disease models will reveal differences in RNA metabolism that underlie cellular heterogeneity and disease progression. While we have used Sp-Cas9 to induce gene knockouts, alternative effectors like base editors and prime editors can be readily incorporated into our modular workflow to identify the role of specific protein domains or regulatory elements on RNA metabolism at nucleotide level resolution. Using non-coding, viral, and synthetic RNAs instead of mRNA reporters has the potential to unlock novel RNA regulatory mechanisms and therapeutic strategies. Finally, expanding ReLiC from our RNA interactome-focused library to all protein coding genes in the human genome will illuminate new interactions between RNA metabolism and other cellular processes.

## Supporting information

Supplemental Tables 1-8 in CSV format, zipped together

## Author Contributions

P.J.N. designed research, performed experiments, analyzed data, and wrote the manuscript. H.P. performed experiments. C.L.W. and A.C.H. assisted with polysome fractionation experiments. C.B., G.Q., and K.Y.C. performed gene ontology analyses. A.R.S. conceived the project, designed research, analyzed data, wrote the manuscript, supervised the project, and acquired funding.

## Acknowledgements

We thank members of the Subramaniam lab, the Basic Sciences Division, and the Computational Biology Program at Fred Hutch for assistance with the project and discussions and feedback on the manuscript. The computations described here were performed on the Fred Hutchinson Cancer Center computing cluster. This research was funded by NIH R35 GM119835 (A.R.S.), NSF MCB 1846521 (A.R.S.), NIH T32 GM008268 (P.J.N.), NIH R37 CA230617 (A.C.H.), NIH R01 CA276308 (A.C.H.), and NIH GM135362 (A.C.H.). This research was supported by the Genomics and Flow Cytometry Shared Resources of the Fred Hutch/University of Washington Cancer Consortium (P30 CA015704) and Fred Hutch Scientific Computing (NIH grants S10-OD-020069 and S10-OD-028685). The funders had no role in study design, data collection and analysis, decision to publish, or preparation of the manuscript.

## Data, Code, and Material Availability

All high throughput sequencing data are publicly available in the NCBI SRA database under BioProject PRJNA1059490. SRA accession numbers with sample annotations are provided as supplementary table S5. All software used in this study are publicly available as Docker images at https://github.com/orgs/rasilab/packages. All other data and analysis code are publicly available at https://github.com/rasilab/nugent_2024. Materials and clarifications pertaining to this study can be publicly requested at https://github.com/rasilab/nugent_2024/issues/new/choose.

## Materials and Methods

### Plasmid construction

Plasmids, oligonucleotides, and cell lines used in this study are listed in supplemental tables S2-S4. DNA sequences of plasmids used in this study are available at https://github.com/rasilab/nugent_2024. Unless specified below, DNA fragments used for cloning were either excised out by restriction digestion or amplified by PCR from suitable templates. Fragments were assembled together using Gibson assembly^100^, and transformed into NEB10beta cells. All constructs were verified by restriction digestion and Sanger or long read sequencing.

### Landing pad vector construction

The *attP* landing pad vector (pHPHS232) was created by using the plasmid backbone, *AAVS1* homology arms, and cHS4 insulator from pASHS11 (pAAVS1P-iCAG.copGFP^101^/Addgene 66577); the Tet-responsive promoter, *attP*, mTagBFP2, P2A, iCasp9, T2A, blasticidin S deaminase, and pCMV-rTTA from pHPHS111 (Addgene 200630); NeoR from pHPHS27 (mtk8b_LA_AAVS1_SA_neoR^102^ /Addgene 123742); and SV40pA from pHPHS5 (mtk4b_002_tSV40^102^ / Addgene 123843).

The *attP\** landing pad vector with *Cas9* (pHPHS800) was created using the plasmid backbone from pYTK089^103^ (Addgene 65196); the cHS4 insulator from pASHS11 (Addgene 66577); the EF1α promoter from pHPHS3 (MTK2_007_pEF1α^102^ /Addgene 123702); *attP\** encoded on oAS1848; *attB* encoded on oAS1482/oAS1540; SpCas9-NLS-FLAG from lentiCas9-Blast^104^ (Addgene 52962); T2A from pPBHS126 (pRRL U6-empty-gRNA-MND-Cas9-t2A-Blast^105^); Hygromycin phosphotransferase (HPH) from pHPHS7 (MTK6_009 CMV-Hygro-bgPA^102^ /Addgene 123863); and SV40pA from pHPHS5 (Addgene 123843).

### Reporter plasmid construction

A base vector for reporter cloning (pHPHS806) was created using the plasmid backbone from pYTK089 (Addgene 65196); the cHS4 insulator, TRE3GV promoter, and T2A-PuroR from pASHS11 (pAAVS1P-iCAG.copGFP/Addgene 66577); EYFP-bGHpA from pPBHS285^106^; *attB\** encoded on oAS1853/oAS1854; mCherry from pHPHS109 (Addgene 171598); and SV40pA from pHPHS5 (Addgene 123843).

Next, reporters were cloned sequentially into the pHPHS806 base vector. We wanted to add unique 6xN barcodes in the 3′ UTR of all reporters to enable sample pooling and multiplexing during sequencing. First, we cloned a series of reporters with unique barcodes in the 3′UTR of the mCherry-puro reporters. A region of the PuroR cassette was digested out of pHPHS806 using BamHI/BsmBI and replaced by Gibson assembly with the same region of PuroR amplified from pHPHS806 using oAS1292 and one of oAS1883-1886, each of which adds a unique 6xN barcode. The resulting plasmids were referred to as pHPHS843-846.

pHPHS843-846 were used as backbones to clone the reporters of interest used for ReLiC screens after digesting out EYFP with KpnI/AgeI and inserting EYFP with a 6xN barcode or β-globin Norm and Ter 39 reporters^36^ (referred to as NTC and PTC here). A 3xFLAG tag was included upstream of the NTC and PTC reporters. The resulting plasmids were referred to as pHPHS853, pHPHS926, and pHPHS927.

### Plasmid library construction

First, a base vector for sgRNA cloning (pHPHS309) was created using the plasmid backbone from pYTK090^103^ (Addgene 65197), amplified using oAS1411/oAS1315; SV40pA from pHPHS5 (mtk4b_002_tSV40^102^/Addgene 123843), amplified with oAS1331/oAS1571; U6 promoter and gRNA scaffold from pAS70 (Brunello library in lentiGuide-puro backbone^31^/Addgene 73178), amplified using oAS1406/oAS1386 and oAS1334/oAS1572, respectively; a GFP dropout cassette from pYTK001^103^ (Addgene 65108), amplified using oAS1407/oAS1408; and a cassette encoding EcoRV and AscI restriction sites, an Illumina R1 sequencing primer binding site, and a T7 promoter, amplified using oAS1573/oAS1574/oAS1577/oAS1578. The R1 primer binding and T7 sequences are for sequencing of sgRNA inserts at the EcoRV site and for *in vitro* transcription from genomic DNA, respectively; the AscI site allows for insertion of reporters and barcodes.

Next, the GFP dropout cassette was excised from pHPHS309 by restriction digestion with BamHI/XhoI and replaced with the custom RBP-targeting dual sgRNA library, which was synthesized by IDT as an oligo pool oAS1899 (Supplementary Table S1) then amplified using oAS1612/oAS1613. Assembled plasmid pools were transformed with high efficiency into NEB10Beta *E. coli* and referred to as pHPHS928.

The reporter barcodes were subsequently added to the sgRNA library plasmid by Gibson assembly using the plasmid backbone from pHPHS309, amplified using oAS1315/oAS1331; the RBP dual sgRNA library from pHPHS928, amplified using oAS1406/oAS1572; and a pair of 20xN barcode sequences, amplified using oAS1573/oAS1574/oAS1575/oAS1576. Assembled plasmid pools were again transformed with high efficiency into NEB10Beta *E. coli*, bottlenecked to ∼5×10^5^ barcode pairs, and referred to as pHPHS932.

Next, an AmpR cassette was inserted between the two sgRNAs in a two-step process. First, an AmpR vector (pHPHS841) was created by Gibson assembly using the plasmid backbone from pYTK083^103^ (Addgene 65190), amplified using oAS1875/oAS1876; AmpR from pYTK083, amplified using oAS1877/oTB11; and the mU6 promoter and tracRNAv2 separated by a HindIII site that was ordered as IDT gBlock oAS1878 and digested with HindIII. The dual sgRNA library in pHPHS932 has two BsmBI restriction sites in between the two sgRNAs that yield sticky ends that are compatible with those generated from the BsaI and NcoI sites flanking the AmpR cassette in pHPHS841. So, the AmpR cassette was then digested out of pHPHS841 using BsaI/NcoI and ligated into BsmBI-digested pHPHS932 library using T4 DNA ligase (Thermo). Ligated plasmid pools were again transformed with high efficiency into NEB10Beta *E. coli* and referred to as pHPHS934.

Next, the mCherry-puro reporters with unique 6xN barcodes were inserted into the pHPHS934 plasmid pool. pHPHS934 was used as the plasmid backbone and was digested with AscI, which cuts between the 20xN barcodes. Barcoded mCherry-puro reporters were digested out of pHPHS853, pHPHS926, and pHPHS927 using NotI, which includes sequence fragments upstream and downstream of the reporters that are homologous to the free ends of AscI-digested pHPHS934 for Gibson assembly. Library diversity was maintained by transformation into high efficiency into NEB10Beta *E. coli* and the resulting plasmids were referred to as pHPHS937, pHPHS938, and pHPHS940.

Finally, the EYFP, β-globin PTC, and β-globin NTC reporters were inserted between the sgRNA cassette and the upstream 20xN barcode sequence in the pHPHS937, pHPHS938, and pHPHS940 libraries with mCherry-puro reporters. pHPHS937-940 were digested with NotI, which cuts immediately upstream of the upstream 20xN barcode sequence. Digesting pHPHS853, pHPHS926, and pHPHS927 with NotI also cut out their EYFP, β-globin PTC, and β-globin NTC reporters flanked by homologous sequences to the free ends of NotI-digested pHPHS937, pHPHS938, and pHPHS940. So, the digested reporters were directly incorporated into pHPHS937, pHPHS938, and pHPHS940 by Gibson assembly. Library diversity was maintained by transformation into high efficiency into NEB10Beta *E. coli* and the resulting plasmids were referred to as pHPHS951, pAS243, and pAS244.

The above sequence of steps to create the final ReLiC libraries is shown schematically in Fig. S1A.

### Lentiviral sgRNA expression plasmid construction

A vector expressing dual sgRNAs targeting *GCN2* was created using the plasmid backbone from pHPHS714 (pJRH051^107^/Addgene 171625), digested with BsmBI; and *GCN2*-targeting sgRNAs encoded on oAS2037/oAS2038 that were PCR amplified to flank the dual sgRNA scaffold using pHPHS928 as a template. The resulting plasmid was referred to as pAS194.

### sgRNA expression plasmid construction for stable integration

A base vector for cloning (pHPHS859) was prepared by Gibson assembly using pHPHS309 as a backbone and bGHpA-*attB\**-mCherry-T2A-puro from pHPHS809 as an insert. sgRNAs targeting *AQR*, *SF3B5*, *SF3B6*, *GCN1* and *FLuc* encoded on oAS2137/oAS2138, oAS2155/oAS2156, oAS2157/oAS2158, oAS2069/oAS2070, and oPN748/oPN749, respectively, were PCR amplified to flank the dual sgRNA scaffold using pHPHS928 as a template. These dual sgRNA PCR products were inserted between the BamHI and XhoI sites of pHPHS859 by Gibson assembly to make pAS298, pAS307, pAS308, pAS232, and pHPHS913.

For RNA-seq of splicing factors, the β-globin reporter from pHPHS927 was inserted into pAS298, pAS307, pAS308, and pHPHS913 by restricting all plasmids with NotI, which generates complementary homology arms that were joined together by Gibson assembly to make pAS310, pAS319-321. For RNA-seq during HHT treatment, the EYFP reporter from pHPHS853 was inserted into pAS232 and pHPHS913 after cutting all plasmids with NotI and joined together by Gibson assembly to make pAS251 and pAS254.

### Cell culture

HEK293T cells (RRID:CVCL_0063, ATCC CRL-3216) were cultured in Dulbecco′s modified Eagle medium (DMEM 1X, with 4.5 g/L D-glucose, + L-glutamine, - sodium pyruvate, Gibco 11965-092) supplemented with 10% FBS (Thermo 26140079) and passaged using 0.25% trypsin in EDTA (Gibco 25200-056). Cells were grown at 37C in 5% CO2. Cell lines were periodically confirmed to be free of mycoplasma contamination.

### Generation of landing pad cell lines

To generate an initial *attP* landing pad line, HEK293T cells were transfected with landing pad plasmid (pHPHS232) and pASHS29 (AAVS1 T2 CRISPR in pX330^108^/Addgene 72833) using polyethylenimine. Cells were selected with 10 μg/ml Blasticidin S, added 96 hours post-transfection. Blasticidin selection was removed after 4 days, and BFP expression was induced by adding 2 μg/ml doxycycline. 24 hours after doxycycline induction, the culture was further enriched for BFP+ cells using a FACSAria II flow cytometer (BD). Clones were isolated by limiting dilution into 96-well plates. After isolating clones, two were pooled into a single cell line (hsPB126).

To integrate a *Cas9* expression cassette with an orthogonal *attP\** site into the initial *attP* landing pad clonal lines, hsPB126 was transfected with Cas9 landing pad plasmid (pHPHS800) and Bxb1 expression plasmid (pHPHS115) using TransIT-LT1 reagent (Mirus). 72 hours post-transfection, hygromycin phosphotransferase (HPH) was induced by adding 2 μg/ml doxycycline, then cells were selected with 150 μg/ml Hygromycin B, added 96 hours post-transfection. After 7 days, doxycycline and Hygromycin B were removed from cells and replaced with 10 μg/ml Blasticidin. Blasticidin selection was ended after 7 days, and this polyclonal cell line (hsPN266) was used for subsequent experiments.

### Integration of plasmid libraries into landing pad

hsPN266 (HEK293T *attP* Cas9*) cells were seeded to 60% confluency in one 15 cm dish per library. 20 μg of *attB\**-containing reporter library plasmid (pAS243, pAS244, pHPHS951) and 5 μg of Bxb1 expression vector (pHPHS115) were transfected per 15 cm dish using TransIT-LT1 reagent (Mirus). Each library was transfected into a single 15 cm dish then expanded into four 15 cm dishes 48 hours post-transfection. Cells were selected with 2 μg/ml puromycin, added 72 hours post-transfection. Puromycin selection was ended after 4 days, and library cell lines (referred to as hsPN305, hsPN306, hsPN285) were contracted back into a single 15 cm dish. 24h after ending puromycin selection, 2 μg/ml doxycycline was added to induce Cas9 expression, and libraries were expanded into three 15 cm dishes – one each for RNA and gDNA harvests the next day plus a third for continued propagation. This splitting procedure was repeated every other day from the propagation dish, so harvests could be taking throughout the duration of the screen. At no point were cultures bottlenecked to fewer than 5×10^6^ cells.

### Library genomic DNA extraction

For each harvest, reporter library genomic DNA was harvested from one 50% confluent 15 cm dish of cells stably expressing the ReLiC library. Genomic DNA was harvested using Quick-DNA Miniprep kit (Zymo), following the manufacturer’s instructions, with 2.5 ml of genomic DNA lysis buffer per 15 cm plate. 30 µg of purified genomic DNA from each library sample was sheared into ∼350 nucleotide length fragments by sonication for 10 minutes on ice using a Diagenode Bioruptor. Sheared gDNA was then *in vitro* transcribed into RNA (denoted gRNA below and in analysis code) starting from the T7 promoter region in the insert cassette using the HiScribe T7 High Yield RNA Synthesis Kit (NEB). Transcribed gRNA was cleaned using the RNA Clean and Concentrator kit (Zymo).

### Library mRNA extraction

For each harvest, reporter library mRNA was harvested from one 50-75% confluent 15 cm dish of cells stably expressing the ReLiC library. Total RNA was harvested by using 3.5 ml of Trizol reagent (Thermo) to lyse cells directly on the plate, and then RNA was extracted from these lysates using the Direct-zol RNA Miniprep kit (Zymo) following the manufacturer’s protocol. polyA+ mRNA was extracted from total RNA using oligo dT25 magnetic beads (NEB). 30-50 μg of total RNA was used as polyA selection input for total barcode counting libraries from each sample while 10-12 μg was used as input for splicing or polysome fraction barcode counting libraries. 4 μl of oligo dT25 beads were used per 1 μg of total RNA input.

### mRNA and genomic DNA barcode sequencing

100-500 ng of polyA-selected mRNA or *in vitro* transcribed gRNA from each library was reverse transcribed into cDNA using SuperScript IV reverse transcriptase (Thermo) following the manufacturer’s protocol. For RT, we used a primer that binds downstream of the 20xN reporter barcode: either oPN777 for mRNA barcode 1, oPN731 for gRNA barcode 1, or oPN779 for mRNA barcode 2. oPN777 and oPN779 contain a 7 nt UMI. Libraries for sequencing total levels of barcode 1 or barcode 2 in each sample were performed in a single step. For both barcodes, a 100-200 μl PCR was performed using Phusion polymerase (Thermo) for 20-25 cycles with cDNA template comprising 1/5th of the final volume, and oPN776 was used as a constant reverse primer that binds the Illumina P5 sequence present on oPN777 and oPN779. Indexed forward primers that bind a constant region upstream of each barcode were used to enable pooled sequencing of different samples (one of oPN730, oPN738, oPN809, oPN815-822, or oJY1-14 for Barcode 1 or one of oPN734, oPN739, or oPN823-825 for Barcode 2). All of these reactions generated a 181 bp amplicon that was cut out from a 2% agarose gel and cleaned using the Zymoclean Gel DNA Recovery Kit (Zymo).

For splicing screens, two rounds of PCR were performed. Round 1 was performed as a 50 μl PCR for 30 cycles, again with cDNA template comprising 1/5th of the final volume and oPN776 as a constant reverse primer. The forward primer for Round 1 was chosen based on the measured splicing event: oPN841 for intron 1 retention, oPN789 for intron 2 retention, or oAS2029 for exon 2 skipping. These generate 532, 302, and 286 bp amplicons, respectively, which were cut out from a 2% agarose gel and cleaned using the Zymoclean Gel DNA Recovery Kit (Zymo), eluting in 15 μl. Round 2 PCR was then essentially the same as the single-step PCR for total Barcode 1 sequencing, except reactions were 20 μl, used 4 μl of cleaned Round 1 product as template, and proceeded for 5 cycles.

Libraries were sequenced on an Illumina NextSeq 2000 using custom sequencing primers. Custom primers for Barcode 1 were oAS1701 for Read 1, oPN732 for Index 1, oPN775 for Index 2, and oPN731 for Read 2. Custom primers for Barcode 2 were oPN735 for Read 1, oPN737 for Index 1, oPN778 for Index 2, and oPN736 for Read 2. Read lengths varied between sequencing runs with 10% phiX spiked in.

### sgRNA insert-barcode linkage sequencing

sgRNA insert-barcode linkages were determined at the step right after barcodes were added to the cloned sgRNA plasmid pool, prior to adding AmpR between the sgRNAs. A 422 bp amplicon containing both sgRNAs and 20xN barcodes was generated from 1.5 ng of pHPHS932 plasmid by 10 cycles of PCR using oKC196/oPN726 primers and Phusion polymerase (Thermo). This product cut out from a 1.5% agarose gel and cleaned using the Zymoclean Gel DNA Recovery Kit (Zymo). This sample was sequenced on an Illumina NextSeq 2000 using custom sequencing primers: oAS1701 for Read 1 (26 cycles), oKC186 for Index 1 (6 cycles), oAS1702 for Index 2 (20 cycles), and oKC185 for Read 2 (75 cycles).

### CRISPR-Cas9 mediated GCN2 knockout for modifier screen

HEK293T cells were seeded to 60% confluency in a 10 cm dish. Cells were transfected with 5 μg of lentiviral transfer plasmid encoding sgRNA targeting *GCN2* (pAS194) or a nontargeting control (pHPHS714), 4 μg of psPAX2 (Addgene #12260), and 1 μg of pCMV-VSV-G (Addgene #8454) using Lipofectamine 3000 reagent (Thermo). Virus was harvested 48 h post-transfection, filtered using a 0.45 micron syringe filter (Genesee), and immediately used to transduce hsPN283 cells that were seeded to 25% confluency in a 15 cm dish. doxycycline was added to 2 μg/ml at the same time as transduction to induce *Cas9* expression, and this culture was maintained from this point and harvested as described in “Integration of plasmid libraries into landing pad”.

### CRISPR-Cas9 mediated gene knockout for RNA-seq

hsPN266 (HEK293T *attP* Cas9*) cells were seeded to 80% confluency in a 6-well dish. 1.6 μg of *attB\**-containing dual sgRNA + reporter plasmid (pAS251,254,310,319-321) and 400 ng of Bxb1 expression vector (pHPHS115) were transfected per well using Lipofectamine 3000 reagent (Thermo). Each construct was transfected into a single well of the 6-well dish then expanded into a 10 cm dish 48 hours post-transfection. Cells were selected with 2 μg/ml puromycin, added 72 hours post-transfection. Puromycin selection was ended after 4 days on these cell lines (referred to as hsAS103,112-114,309,313). After ending puromycin selection, 2 μg/ml doxycycline was added to induce Cas9 expression. Cells were grown in 6-well plates in the presence of doxycycline for 4 days then harvested for RNA-seq.

### Polysome profiling

After Cas9 induction, 293T cells expressing ReLiC libraries were passaged for 6 days. On day 6, lysates were prepared from each library at 30% confluency in a 15 cm dish. Cultures were treated with 100 μg/ml cycloheximide for 5 minutes prior to harvest, then cells were trypsinized (including 100 μg/ml cycloheximide) and pelleted at 300xg for 5 min. Cell pellets were lysed on ice in 300 μl of polysome lysis buffer (10 mM Tris-HCl pH 7.4 (Ambion), 132 mM NaCl (Ambion), 1.4 mM MgCl2 (Ambion), 19 mM DTT (Sigma), 142 μg/ml cycloheximide (Sigma), 0.1% Triton X-100 (Fisher), 0.2% NP-40 (Pierce), 607 U/ml SUPERase-In RNase Inhibitor (Invitrogen)) with periodic vortex mixing. Lysates were clarified by centrifugation at 9300xg for 5 min and supernatants were transferred to fresh tubes. This total lysate was split into two parts: 50 μl for total mRNA isolation, and 250 μl for polysome profiling. For each sample, the 250 μL lysate fraction was layered onto a 10%–50% (w/v) linear sucrose gradient (Fisher) containing 2 mM DTT (Sigma) and 100 μg/mL heparin (Sigma). The gradients were centrifuged at 37,000 rpm for 2.5 h at 4°C in a Beckman SW41Ti rotor in Seton 7030 ultracentrifuge tubes. After centrifugation, samples were fractionated using a Biocomp Gradient Station by upward displacement into collection tubes, through a Bio-Rad EM-1 UV monitor (Bio-Rad) for continuous measurement of the absorbance at 260 nm. 820 μl of TRIzol Reagent (Invitrogen) were added to each RNA fraction. Total (input), monosome-associated (fraction 4 and 5), low polysome-associated (fractions 6-9), and high polysome-associated (fractions 10-13) mRNA samples were isolated from TRIzol (Invitrogen) using the Direct-zol RNA Miniprep Plus Kit (Zymo Research) with DNaseI treatment according to manufacturer’s directions.

To examine whether GCN1 affects the level of RNAse-resistant disomes during HHT treatment, polysome profiling was performed with four different samples: 293T cells expressing sgGCN1 and sgFLUC from “CRISPR-Cas9 mediated gene knockout for RNA-seq” after 1 week of Cas9 induction, treated for 1 hour with 1 μM HHT or DMSO. Polysome profiling was performed similar to the Polysome ReLiC screen, but with the following modifications. Each sample was harvested from one 10-cm dish at 70% confluency. Prior to loading onto sucrose gradients, lysates were incubated with or without the addition of 1 U of micrococcal nuclease per μg of RNA and 5 μM CaCl_2_ at room temperature for 1 hour. Micrococcal nuclease digests were quenched by addition of 5 μM EGTA prior to loading on sucrose gradients.

### RNA-seq

RNA was isolated using the Direct-zol RNA Miniprep kit (Zymo). Sequencing libraries were generated with the NEBNext Ultra II Directional RNA Library Prep Kit (NEB) and sequenced on a NextSeq 2000 (Illumina) with 2×50 cycle paired-end reads.

### Ribosome profiling

Ribosome profiling was performed with four different samples: 293T cells expressing sgGCN1 and sgFLUC from “CRISPR-Cas9 mediated gene knockout for RNA-seq” after 1 week of Cas9 induction, treated for 1 hour with 1 μM HHT or DMSO. For each sample, we used one 15-cm plate of cells, seeded to ∼40% confluence at harvest. Ribosome profiling protocol was adapted from^109^ with the following modifications. For sample harvesting, we removed media from each plate and flash froze samples by placing the plate in liquid nitrogen and transferred to −80 °C until lysis. We performed nuclease footprinting treatment by adding 80 U RNase I (Invitrogen AM2294) to 25 μg of RNA. We gel-purified ribosome protected fragments with length between 26 and 34 nucleotides using RNA oligo size markers. We used polyA tailing instead of linker ligation following previous studies^110,111^. Libraries were sequenced on an Illumina Nextseq 2000 in 50bp single end mode.

### Immunoblot analysis

sgGCN1 and sgFLUC cell lines used for RNA-seq were incubated with drugs at indicated concentrations for 30 or 60 min before harvest. Homoharringtonine (Biosynth, FH15974) and anisomycin (RPI, A50100) were dissolved in DMSO. Cells were rinsed with PBS and lysed in RIPA buffer. Lysates were kept on ice during preparation and clarified by centrifugation at 15,000 rpm for 10 min. After clarification, supernatants were boiled in Laemmli loading buffer containing DTT, and Western blots were performed using standard molecular biology procedures Proteins were resolved by 4%–20% Criterion TGX protein gels (Bio-Rad) and transferred to PVDF membranes using a Trans-Blot Turbo transfer system (Bio-Rad). Membranes were blocked with 5% BSA (Thermo) in TBST and incubated with primary antibodies overnight at 4°C with gentle rocking. Blots were washed with TBST, then incubated with secondary antibodies diluted in TBST + 5% BSA for 1 hr at RT with gentle rocking. Membranes were washed again with TBST, developed using SuperSignal West Femto Maximum Sensitivity Substrate (Thermo), and imaged on a ChemiDoc MP imaging system (Bio-Rad).

### Flow cytometry

After dissociating cells from culture dishes, they were pelleted and resuspended in Dulbecco’s phosphate-buffered saline (Gibco 14190-144) supplemented with 5% FBS. Forward scatter (FSC), side scatter (SSC), BFP fluorescence (BV421), YFP fluorescence (FITC), and mCherry fluorescence (PE.Texas.Red) were measured for 10,000 cells in each sample using a BD FACS Symphony or Fortessa instrument.

### qRT-PCR

Plasmids that express the β-globin PTC and NTC reporters along with mCherry-Puro (pHPHS926 and pHPHS927) were integrated into hsPN266 cells by transfection using TransIT-LT1 reagent and pHPHS115 Bxb1 expression plasmid. After Puromycin selection, cells were grown in the presence of 2 μg/ml doxycycline for 4 days then RNA was harvested from both samples from a 6-well plate at 30% confluency. cDNA was prepared from 500 ng of total RNA using random hexamer primers and Maxima RT enzyme (Thermo). cDNA reactions were diluted 1:10, then 4 μl of diluted cDNA were used as template in 20 μl qPCR reactions using Phusion polymerase (Thermo) and SYBR Green (Thermo) run on a QuantStudio5 thermocycler (Thermo). Reactions were performed as 3 biological replicates using oPN719/oPN731 for the β-globin reporters or oPB466/oPB467 for mCherry.

### Computational analyses

Pre-processing steps for high-throughput sequencing were implemented as Snakemake^112^ workflows run within Singularity containers on an HPC cluster. Python (v3.9.15) and R (v4.2.2) programming languages were used for all analyses unless mentioned otherwise. All software used in this study are publicly available as Docker images at https://github.com/orgs/rasilab/packages.

### Barcode to insert assignment

Raw data from insert-barcode linkage sequencing are in FASTQ format. Barcode and sgRNA insert sequences were extracted from corresponding reads and counted using awk; sgRNA inserts and corresponding barcodes were omitted if the sequenced sgRNA insert was not present in the designed sgRNA library (oAS1899). The remaining barcodes were aligned against themselves by first building an index with bowtie2-build with default options and then aligning using bowtie2 with options -L 19 -N 1 --all --norc --no-unal -f. Self-alignment was used to exclude barcodes that are linked to distinct inserts or ones that are linked to the same insert but are aligned against each other by bowtie2 (presumably due to sequencing errors). In the latter case, the barcode with the lower count is discarded in filter_barcodes.ipynb. The final list of insert-barcode pairs with a minimum of 5 reads is written as a comma-delimited .csv file for aligning barcodes from genomic DNA and mRNA sequencing below.

### Barcode counting in genomic DNA and mRNA

Raw data from sequencing barcodes in genomic DNA and mRNA are in FASTQ format. Barcode and UMI sequences were extracted from corresponding reads, counted using awk, and assigned to reporters based on their unique 6xN identifier. Only distinct barcode-UMI combinations where the barcode is present in the filtered barcodes .csv file from linkage sequencing are retained. The number of UMIs per barcode and associated insert are written to a .csv file for subsequent analyses in R. Only barcodes with a minimum of 20 UMIs were used for analysis. Barcode counts from pairs of samples were used to run MAGeCK^38^ with --additional-rra-parameters set to --min-number-goodsgrna 3. sgRNAs without a minimum of 20 UMI in one of the compared samples were set to 20 UMI counts before running MAGeCK.

### RNA-seq analyses

Raw reads were aligned against the human genome (GRCh38) along with transcript annotations from Ensembl (v108, *Homo sapiens*). Only primary chromosomes (1-22, X, MT) were used for sequence alignment and downloaded from https://ftp.ensembl.org/pub/release-108/fasta/homo_sapiens/dna/. Transcript annotations were downloaded from https://ftp.ensembl.org/pub/release-108/gtf/homo_sapiens/Homo_sapiens.GRCh38.108.gtf.gz, and subset using awk to include only transcripts on primary chromosomes. Reference index for alignment using STAR v2.7.11a with options --runThreadN: 36, --runMode: genomeGenerate, --sjdbGTFfile: gtf file from above, --limitSjdbInsertNsj: 3000000, --genomeFastaFiles fasta files from above. Alignment was performed using STAR with options --runThreadN: 36, --runMode: alignReads, --alignSJoverhangMin: 300, --alignSJDBoverhangMin: 6, --outSAMmultNmax: 1, --quantMode: GeneCounts, --readFilesCommand: zcat. All annotated splice junctions were extracted from the GTF file using extract_splice_site_annotations.py which closely followed the script hisat2_extract_splice_sites.py from HISAT2. Start and end coordinates of the spliced-out intron were used to designate splice junctions. Annotated cassette exons were identified as those splice junction coordinates that contain exactly 1 exon within them, exactly 2 introns within them, and either the 5’ or the 3’ end of the enclosed introns being the same as the 5’ or the 3’ end of the parent splice junction. Number of reads aligning to each splice junction was extracted from the STAR output file SJ.out.tab. To quantify exon skipping, we used a percent spliced out metric (100 - percent spliced in) since most of the cassette exons that were skipped were fully included in unperturbed cells. Percent spliced out was calculated using junction reads aligning to the skipped isoform junction (minimum threshold of 2 reads) divided by the sum of junction reads aligning to the skipped isoform junction and the junction reads aligning to the flanking included isoform junctions (minimum threshold of 100 reads summed across the two flanking junctions). Number of reads aligning to each intron was calculated using the alignments file and splice junction annotation file using the GenomicRanges function findOverlaps with a minimum overlap of 10nt and a minimum threshold of 100 reads. The intron read count was normalized to per nt by multiplying by a factor of read_length / (intron_length + read_length) to account for the difference in length between introns. Percent spliced in was calculated as the normalized intron read count divided by the sum of the normalized intron read count and the read count for the annotated splice junction with the intron splice out. Gene counts file from STAR was used to perform differential expression analysis using DESeq2 1.38.0.

### Ribosome profiling analyses

PolyA adapters were trimmed from sequencing reads using cutadapt 4.4 with parameters -a AAAAAAAAAA --minimum-length=22 --match-read-wildcards. Trimmed reads were aligned to ribosomal RNA contaminant sequences (NCBI accession NR_003287.2, NR_003286.3, NR_023363.1, and NR_003285.2) using bowtie 1.3.1 with default parameters. Trimmed reads that did not align to ribosomal RNA were aligned against human transcripts (MANE v1.3) using bowtie 1.3.1 with parameters --norc --no-unal --sam. Aligned reads were converted to BAM format, sorted and indexed using samtools 1.16.1. Aligned reads between 27 nt and 33 nt were trimmed by 13nt from their 5′ end to identify the location of the P-site. The location of the P-site relative to the annotated start codon of each transcript was calculated using the start codon annotation in MANE v1.3. Reads assigned to each location relative to the start codon of all transcripts were summed and normalized by the maximum value across all locations to calculate the metagene ribosome P-site profile. P-site profile for individual genes was calculated by summing reads assigned to each location along the unique MANE transcript for that gene. Analysis of previous ribosome profiling studies was performed using the same pipeline as above with the following modifications. We used the ribosome profiling data from harringtonine- or lactimidomycin-treated samples corresponding to SRA accession numbers SRR1802157, SRR1802156, SRR1802155, SRR1802151, SRR1802150, SRR1802149, SRR1802136, SRR1802135, SRR1802134, SRR1802133, SRR1333394, SRR4293695, SRR4293693, SRR1630828, SRR1630830, SRR1630829, SRR6327777, SRR9113062, SRR9113063, SRR2732970, SRR2954801, SRR2954800. The sequences CTGTAGGCACCATCAAT and AGATCGGAAGAGC were used from adapter trimming. P-site counts across all samples were summed to calculate the profile for individual genes.

### Perturb-seq analyses

Normalized bulk expression profiles from genome-wide Perturb-seq data^16^ were downloaded from Figshare as the file K562_gwps_normalized_bulk_01.h5ad. Data were subset to include only the genes of interest, transcripts with infinite expression were removed, and Pearson correlation coefficients between all pairs of expression profiles were calculated using the R function corr.test.

### Gene ontology analyses

The gene_summary.txt output from MAGeCK was ordered either by pos|fdr for positive fold changes or by neg|fdr for negative fold changes and input into the web interface of GOrilla^113^. Enriched cellular processes and components were manually curated for representative GO terms with minimal overlap of genes.

### Chemicals

| Reagent | Source | Identifier |
| --- | --- | --- |
| ISRIB | Sigma | SML0843 |
| Homoharringtonine | Biosynth | FH15974 |
| Anisomycin | Research Products International | A50100 |
| Hygromycin B | Research Products International | H75020 |
| Puromycin dihydrochloride | Research Products International | P33020 |

### Antibodies

| Reagent | Source | Identifier |
| --- | --- | --- |
| p38 MAPK | Cell Signaling Technology | 8690; RRID:AB_10999090 |
| Phospho-p38 (Thr180/Tyr182) | Biolegend | 690201; RRID:AB_2801132 |
| GCN1 | Bethyl | A301843AT; RRID:AB_1264319 |
| Goat Anti-Rabbit IgG (H L)-HRP Conjugate | Bio-Rad | 1721019; RRID:AB_11125143 |
| Goat Anti-Mouse IgG (H L)-HRP Conjugate | Bio-Rad | 1721011; RRID:AB_11125936 |

## Supplementary Figures

**Figure S1.**
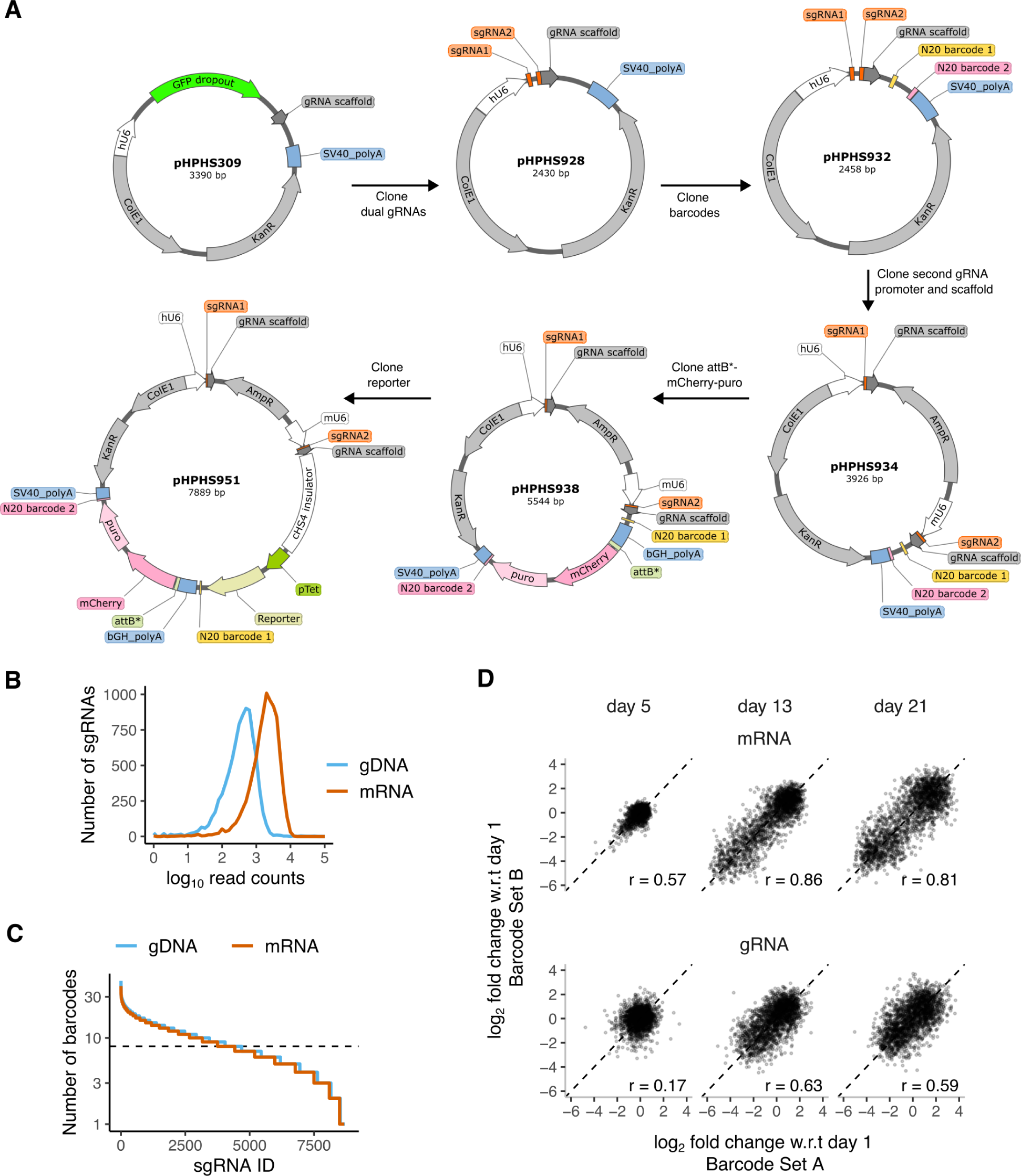
ReLiC library design and validation. **A.** *Depiction of cloning scheme for ReLiC library and reporters.* **B.** *Distribution of barcode read counts for sgRNA pairs in mRNA and genomic DNA*. **C.** *Number of unique barcodes linked to each sgRNA in ReLiC library*. **D.** *Correlation between distinct barcode sets in ReLiC fitness screens*. Each point represents a unique sgRNA pair from the ReLiC RBP library. For each sgRNA pair, individual linked barcodes were randomly partitioned into two sets of equal size (or to within a barcode for odd number of detected barcodes). r refers to Pearson correlation coefficient between the barcode sets.

**Figure S2.**
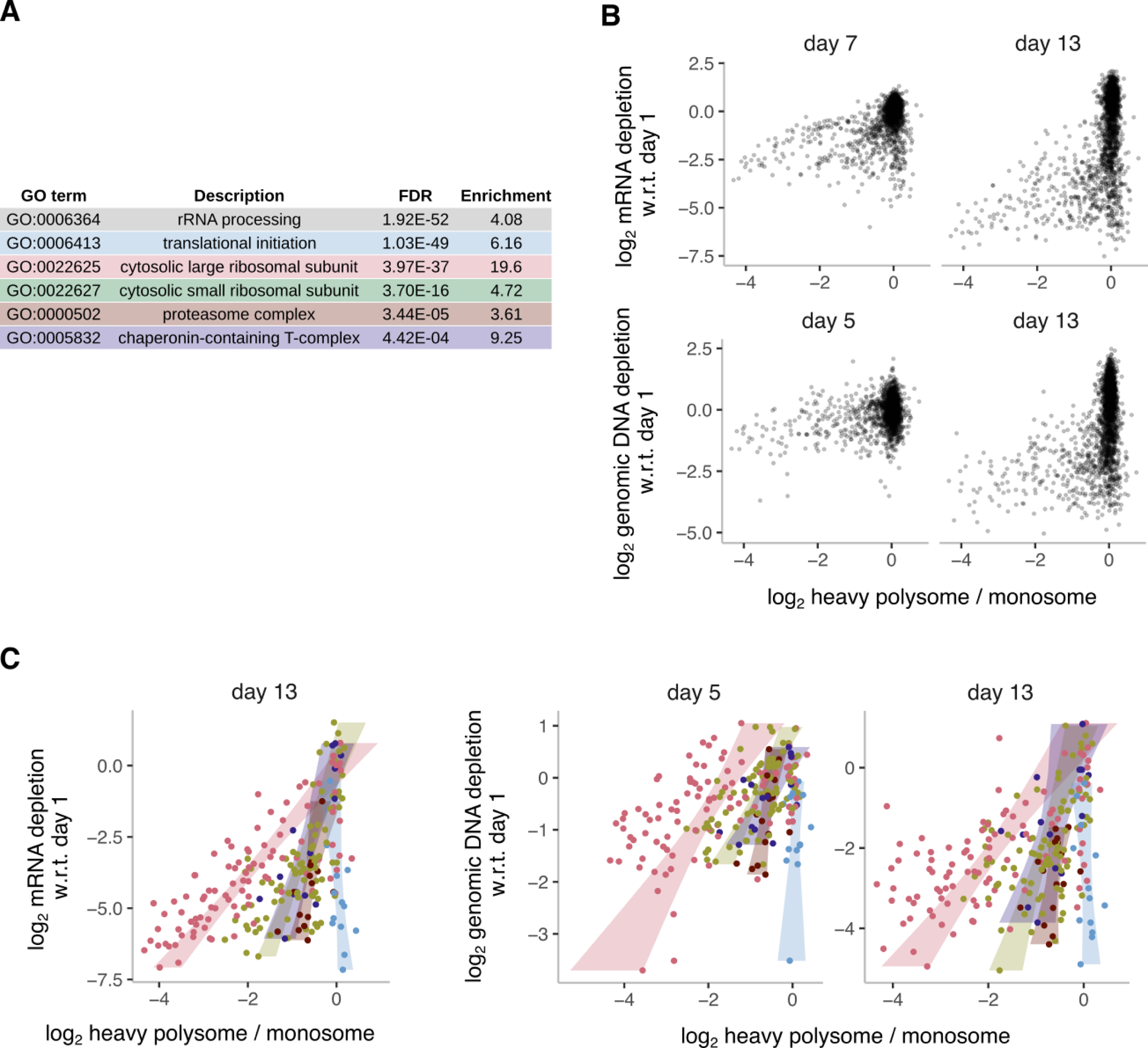
Polysome ReLiC screen for regulators of mRNA translation. **A.** *Gene ontology analysis of perturbations that decrease heavy polysome to monosome ratio.* Gene ontology analysis performed using GOrilla^113^ and a subset of enriched terms representative of specific gene classes are shown. **B.** *Comparison of heavy polysome to monosome ratio with growth fitness measured by mRNA and genomic DNA barcode seqencing 13 days after Cas9 induction for all gene knockouts*. **C.** *Comparison of heavy polysome to monosome ratio with growth fitness measured by genomic DNA barcode sequencing for gene knockouts fcin specific groups.* Points correspond to genes targeted in the ReLiC-RBP library. Shaded areas correspond to 95% confidence intervals for a linear fit of polysome to monosome ratio to growth fitness within each gene group.

**Figure S3.**
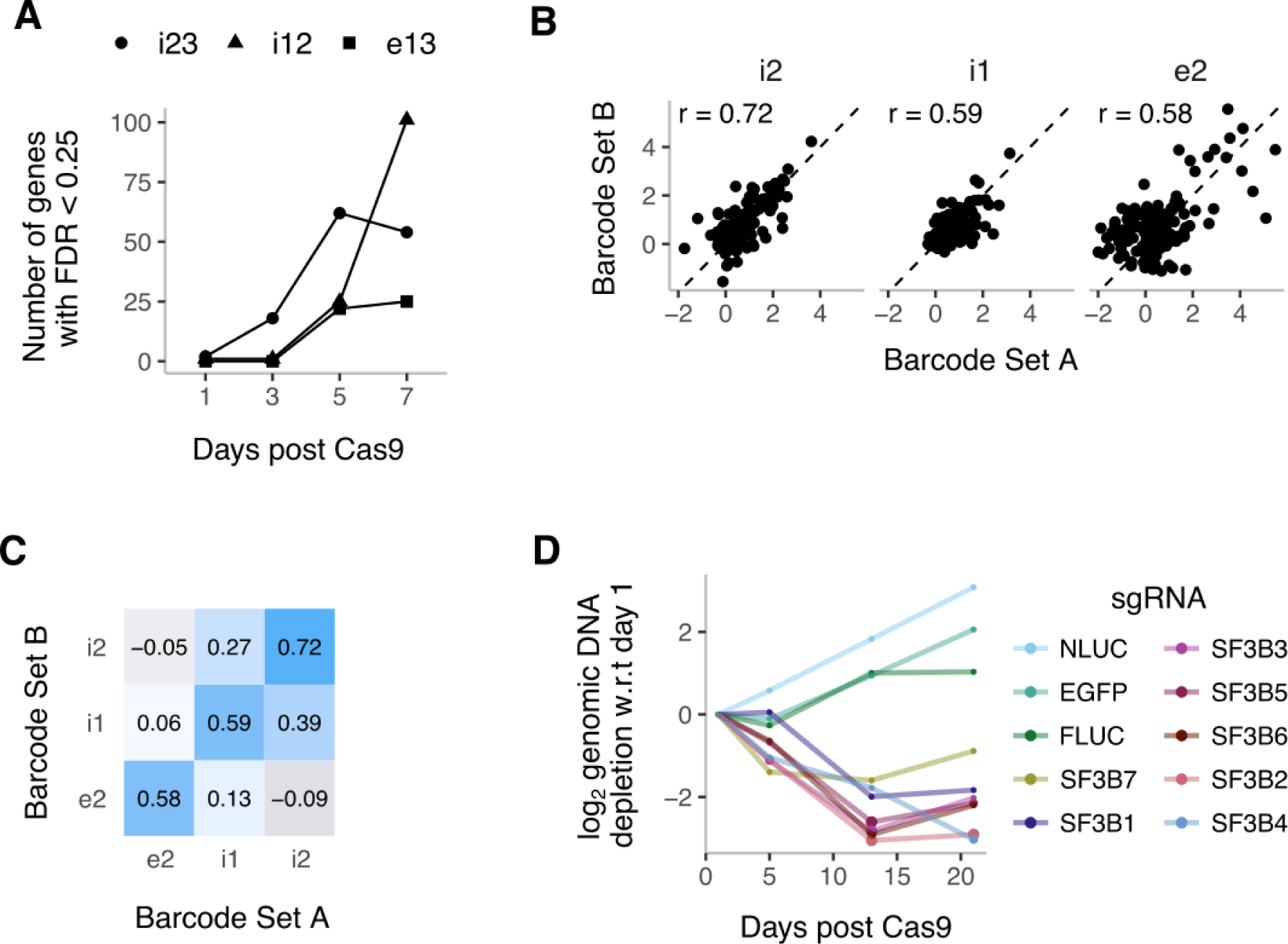
Isoform-specific splicing screen using ReLiC. **A.** *Number of gene hits that increase the level of the indicated reporter isoform on indicated days after Cas9 induction.* **B.** *Correlation between barcode sets*. For each sgRNA, individual linked barcodes were randomly partitioned into two sets, as in Fig. S1D. Each point represents a unique gene that was classified as a hit either with barcode Set A or barcode set B. r refers to Pearson correlation coefficient between barcode sets. **C.** *Correlation between relative levels of different mRNA isoforms*. Values represent Pearson correlation coefficients for pairwise comparison between the two barcode sets in B. **D.** *Depletion of genomic DNA barcodes corresponding to SF3b complex subunits after Cas9 induction*.

**Figure S4.**
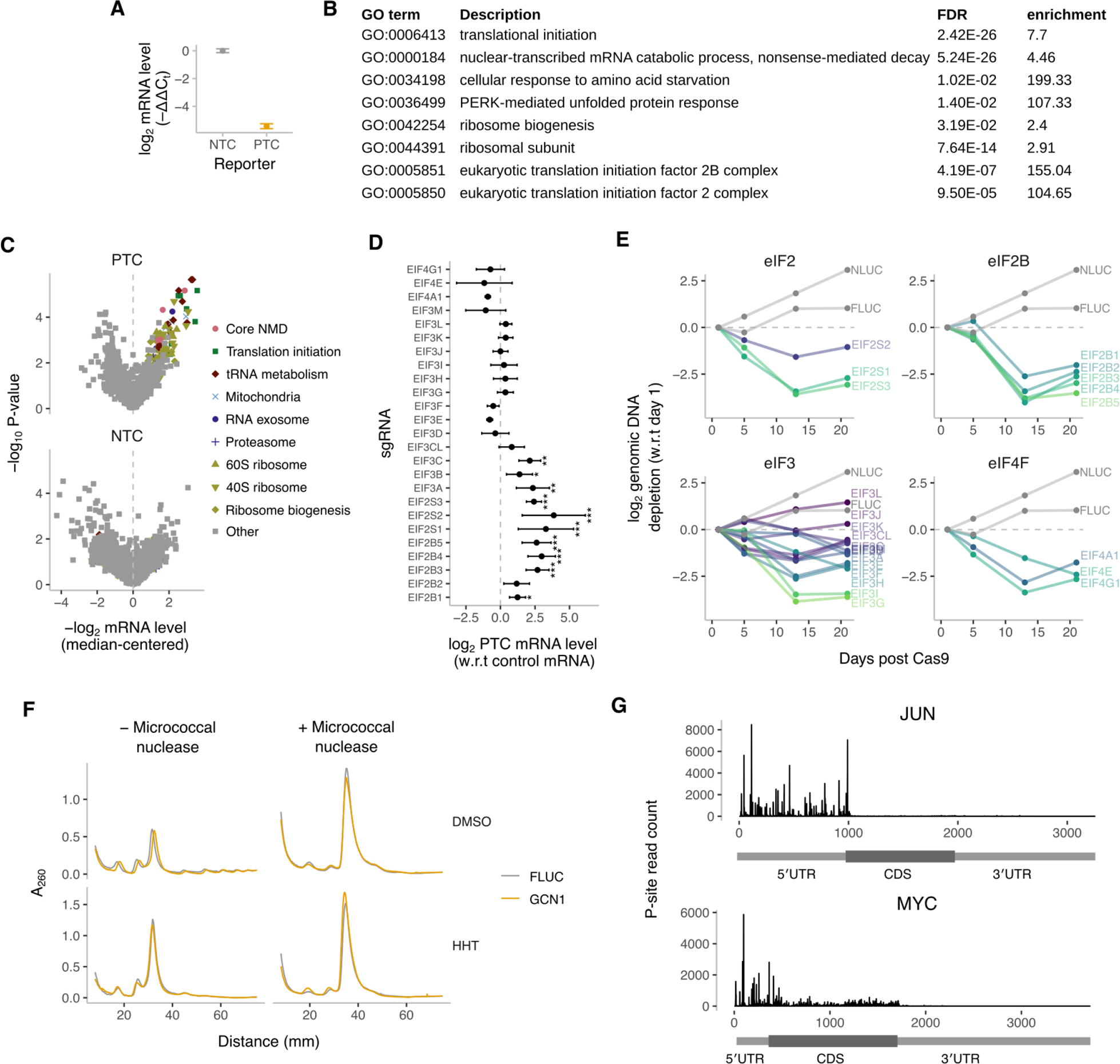
Dissecting mRNA quality control using ReLiC. **A.** *Validation of β-globin NMD reporters.* Relative reporter mRNA levels measured by qPCR (n=3). Y-axis represents -ΔΔC_t_ value of indicated reporter mRNA relative to mCherry-Puro control mRNA. **B.** *Gene ontology analysis of perturbations that increase PTC reporter mRNA levels*. **C.** *Volcano plot of reporter mRNA levels with dual barcode screen*. Each point corresponds to a gene targeted by the ReLiC library. Marker shape and color denotes one of highlighted gene groups. Genes with FDR < 0.05 and belonging to one of the highlighted groups are listed in the legend. **D.** *PTC reporter levels for individual translation initiation complex subunits.* Points denote mean and error bars denote standard deviation across sgRNAs for each gene. P-values are as calculated by MAGeCK. **E.** *Growth fitness after depletion of translation initiation complex subunits*. **F.** *Polysome profiles of GCN1-depleted and control cell lines after HHT treatment*. Cells were treated with 1 μM HHT or DMSO for 1 hour prior to lysis. Polysome lysates were digested with 1 U micrococcal nuclease / μg of RNA prior to sucrose gradient sedimentation to isolate RNAse-resistant monosomes and disomes. **G.** *Ribosome P-site density on JUN and MYC mRNAs from previous ribosome profiling studies using harringtonine or lactimidomycin to arrest initiating ribosomes*.

## Supplementary Table Descriptions

**S1: sgRNA pairs and genes targeted in the ReLiC-RBP library**

**S2: Plasmids used for this study**

**S3: Oligonucleotides used for this study**

**S4: Cell lines used for this study**

**S5: SRA accession numbers**

**S6: Read counts for sgRNAs**

**S7: MAGeCK output for sgRNA comparisons**

**S8: MAGeCK output for gene comparisons**

## Footnotes

1 We refer to ‘polysome to monosome ratio of barcode counts’ as ‘polysome to monosome ratio’ for brevity, but we emphasize the distinction from the polysome to monosome ratio of A_260_ as typically used in the polysome profiling literature.

